# Fast dendritic excitations primarily mediate back-propagation in CA1 pyramidal neurons during behavior

**DOI:** 10.64898/2026.01.03.696606

**Authors:** Byung Hun Lee, Pojeong Park, Xiang Wu, J. David Wong-Campos, Junjie Xu, Marley Xiong, Yitong Qi, Yi-Chieh Huang, Daniel G. Itkis, Sarah E. Plutkis, Luke D. Lavis, Adam E. Cohen

## Abstract

Dendrites integrate synaptic inputs to trigger action potentials, and dendrites carry back-propagating action potentials (bAPs) to synapses where these signals contribute to plasticity. Despite strong evidence for a rich repertoire of nonlinear dendritic excitations, the *in vivo* roles of these excitations in dendritic integration and back-propagation remain uncertain. Here, we used high-speed voltage imaging through a chronically implanted microprism to map membrane potential dynamics from basal to apical dendrites of CA1 neurons in mice navigating in a virtual reality environment. Despite complex dendritic branch morphology, the dynamics were largely captured by 2 or 3 electrical compartments: basal, soma, and apical. Fast dendritic spikes almost always started from bAPs, indicating that dendritic spikes are primarily a consequence rather than a cause of somatic spiking. These fast spikes sometimes triggered slower apical dendritic plateau depolarizations, which drove complex spikes at the soma. We found that the biophysics of dendritic excitability determined the distribution of simple and complex spikes across a place field. Our results show how CA1 pyramidal neurons convert synaptic inputs to spiking outputs and suggest a primary role of dendritic nonlinearities in mediating activity-dependent plasticity.

## Introduction

Dendrites integrate synaptic inputs to drive neural firing, and carry back-propagating action potentials (bAPs) from the cell body to the synapses to mediate plasticity (*1–4*). The diverse ion channels in dendrites may mediate both processes, though the computational requirements of integration and back-propagation are very different. Many studies have observed dendritic nonlinearities and regenerative spikes—including Na^+^ (*5*, *6*), Ca^2+^ (*7*, *8*), and N-methyl-D-aspartate receptor (NMDAR) spikes (*9*) and have characterized the underlying biophysics in acute slices (reviewed in (*3*)). However, it is not clear to what extent dendritic spikes are cause vs. consequence of somatic spikes *in vivo*.

Dynamic excitation/inhibition balance and modulatory tone may lead to very different dendritic dynamics *in vivo* vs. in slices. Prior *in vivo* dendritic patch-clamp recordings (*10–13*) and voltage imaging experiments (*14–16*) only sampled sub-regions of the dendritic tree, and thus did not probe the full spatiotemporal dynamics of dendritic excitations. Here, we applied voltage imaging through a chronically implanted prism (*17*) to map dendritic voltage at kHz frame rates from basal to apical dendrites of CA1 pyramidal neurons in mice navigating in a virtual reality (VR) environment. By combining spatial mapping and targeted perturbations of dendritic voltage dynamics, we studied how dendritic excitations contribute to neural integration and plasticity.

## Results

We expressed the Optopatch-V construct (*18*) in a sparse subset of CA1 pyramidal neurons of adult mice. Optopatch-V comprises the chemogenetic voltage indicator, Voltron2-JF608 (*19*), with the blue-shifted channelrhodopsin, CheRiff (*20*), both tagged with Lucy-Rho motifs (*21*) to improve membrane trafficking in dendrites (**Fig. 1A**). Voltron2 is a negative-going indicator; to avoid confusion, we flipped the sign of the responses so depolarization is always reported as a positive signal. A custom microprism was implanted in the hippocampus along the anterior-posterior axis (*17*), to provide a side-on view of CA1 pyramidal neurons (**Fig. 1B**, **Fig. S1** and **Movie S1**). We used a custom microscope (described in (*16*)) equipped with two digital micromirror devices (DMDs) to independently pattern fluorescence excitation (607 nm) and optogenetic stimulation (488 nm; **Fig. 1C**). The field of view was ∼450 × 210 μm, spanning basal to near the end of the distal apical dendrites, imaged at 1 kHz. Comparison of the functional data with spinning disk confocal structural reconstructions of the whole dendritic tree showed that the recorded dendrites comprised 69 ± 17% of the total dendritic tree (mean ± s.d., *n* = 15 neurons, 10 mice; **Fig. 1D**, **Fig. S2**, and **Movie S2**).

**Figure 1.**
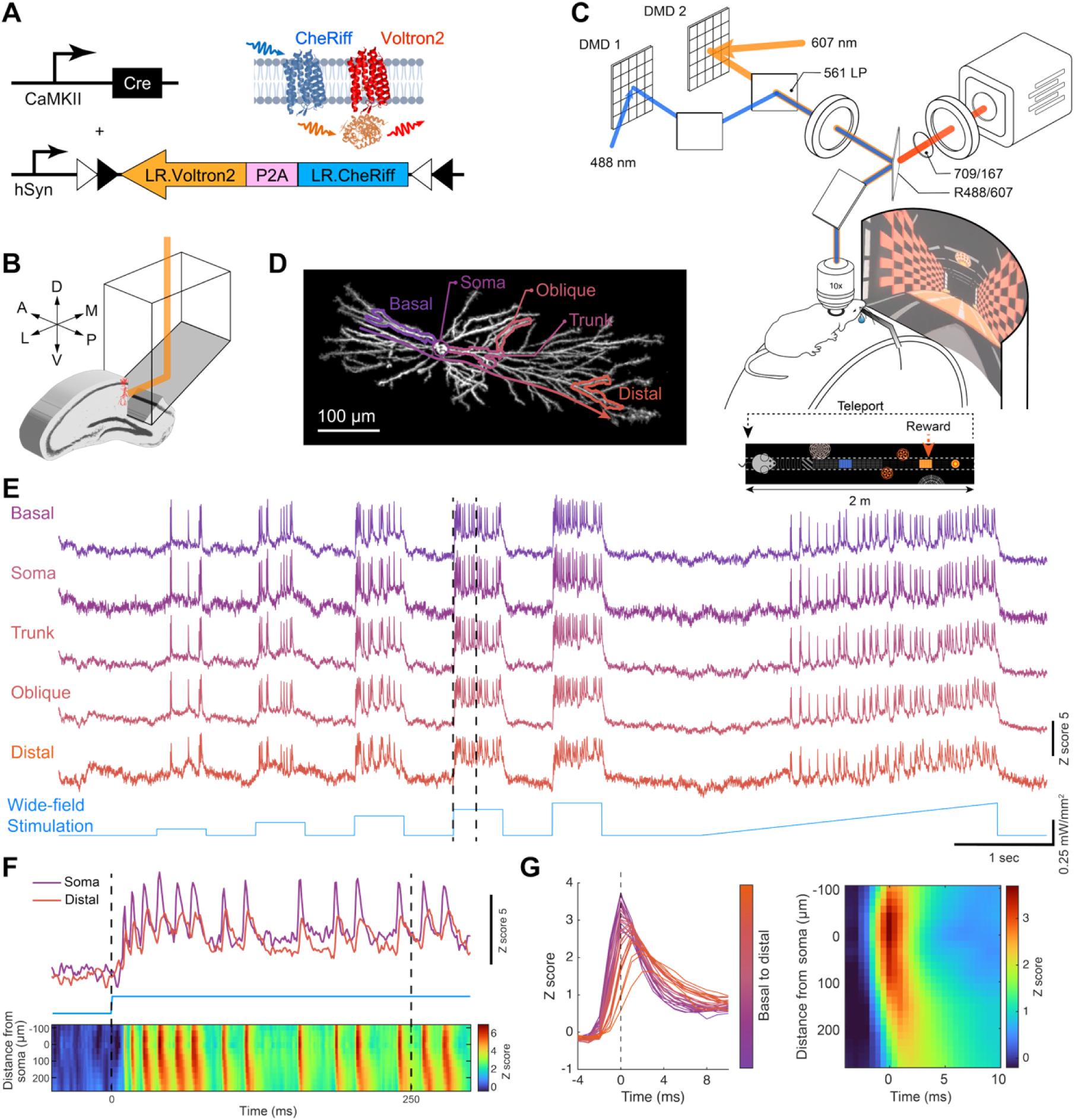
Dendritic voltage imaging of CA1 pyramidal neurons in vivo. (A) Genetic constructs to express Optopatch-V, comprising a voltage indicator, Voltron2-JF608, and optogenetic actuator, CheRiff. Lucy-Rho (LR) motifs were used to improve dendritic trafficking of both optogenetic tools. (B) Microprism (1.5 × 1.5 × 2.5 mm) implanted along the anterior-posterior axis provides a side-on view of CA1 pyramidal neurons. (C) The optical setup contained two digital micromirror devices (DMDs) for targeted illumination and targeted optogenetic stimulation. (D) Maximum z-projection of a spinning disk confocal image of a CA1 pyramidal neuron imaged through a prism in an anesthetized mouse, imaged via Voltron2-JF608 fluorescence. (E) Voltage traces from dendritic branches of the neuron shown in (D) during wide-area optogenetic stimulation in an anesthetized mouse. (F) Zoomed-in trace of the dotted region in (E). Bottom: corresponding voltage kymograph along the basal-apical axis (arrow line in (D)) showing bAP propagation along the apical dendrite. (G) Spike-triggered average (n = 94 spikes) of the spike waveform. Left: color-coded from basal to apical dendrites. Right: kymograph along the basal-apical axis showing spike initiation at the soma, and propagation delays, and attenuation along the dendrites.

To assess the specificity of the targeted optogenetic stimuli, we projected a bar of blue light (26 x 5.8 μm) at a series of transverse offsets from target dendrites in an isoflurane-anesthetized mouse. The optogenetically evoked depolarization decreased to 20% of its peak value for a transverse offset of 13.5 ± 3.9 μm (mean ± s.d., *n* = 6 branches, 3 neurons, 2 mice, **Fig. S3**), confirming that optogenetic stimuli could be targeted with ∼15 μm precision.

To assess the fidelity of branch-specific voltage signals, we calculated spike-triggered average images of the quantity ΔF = F(spike)-F(baseline). The ΔF voltage footprint decreased to 20% for a transverse offset of 9.7 ± 4.7 μm from the dendritic shaft (mean ± s.d., *n* = 15 neurons, 7 mice, **Fig. S3**). These results showed that structured illumination allowed extraction of branch-specific voltage signals.

To extract voltage traces from the recordings, we developed an image-processing pipeline that comprised corrections for camera rolling shutter timing, brain motion, and photobleaching; masking of blood vessels and autofluorescent puncta; and calculation of optimal weight masks to extract voltage signals (**Fig. S4** and **Methods**). To account for spatially varying background and Voltron2 density, we normalized the voltage signal at each dendritic location by a shot-noise-robust estimate of the amplitude of the local voltage fluctuations (**Fig. S5**, **Movie S4** and **Methods**).

Under wide-area optogenetic stimulation in an isoflurane-anesthetized mouse, blue light robustly evoked action potentials in proportion to blue-light intensity (**Fig. 1E-F**). Spike-triggered average movies (triggered off spike timing at the soma) produced high-resolution bAP maps, revealing spike initiation at the soma, and attenuation and delay along both the apical and basal dendrites (**Fig. 1G**). Using the Sub-Nyquist Action Potential Timing (SNAPT) technique (*20*) we inferred the pixel-by-pixel sub-millisecond timing of the bAP wavefront, showing initiation at the soma and back-propagation along the apical and basal dendrites (**Fig. S6** and **Movie S4**).

Mice were head-fixed on a linear track in a 2 m long virtual reality (VR) environment (**Fig. 1C**). During voltage imaging, mice explored a novel environment never presented during the prior 2 weeks of daily training, and received a water reward once per lap (**Methods**). The firing properties of neurons in prism-implanted mice undergoing VR navigation did not differ significantly from neurons observed through a standard hippocampal cannula window (*22*), in terms of firing rate, burst rate, burst size, and spatial information (**Fig. S7A–C**, **F**). In both preparations, optogenetic stimulation evoked plateau potentials and induced formation of place fields whose peak activity occurred slightly ahead of the stimulation position in the VR space. These properties are characteristic of behavior time-scale plasticity (BTSP) (*23–25*) (**Fig. S7D–E**).

### Dendritic subthreshold activity is correlated within lamina

We first investigated dendritic subthreshold dynamics in mice navigating in VR. Reliable voltage imaging was maintained for up to 10 minutes continuously, and mice ran 9–32 laps during this period (**Fig. 2A, B**). Fluorescence brightness after 10 min of continuous voltage imaging was 72 ± 14% of the initial value (mean ± s.d., *n* = 20 neurons, 12 mice). This modest photobleaching was corrected in post-processing. Subthreshold voltage traces were obtained by excluding data around spike times ± 3 ms, linearly interpolating from -4 to +4 ms, and then smoothing with a 20 ms sliding average (**Fig. 2C**, **Methods**).

**Figure 2.**
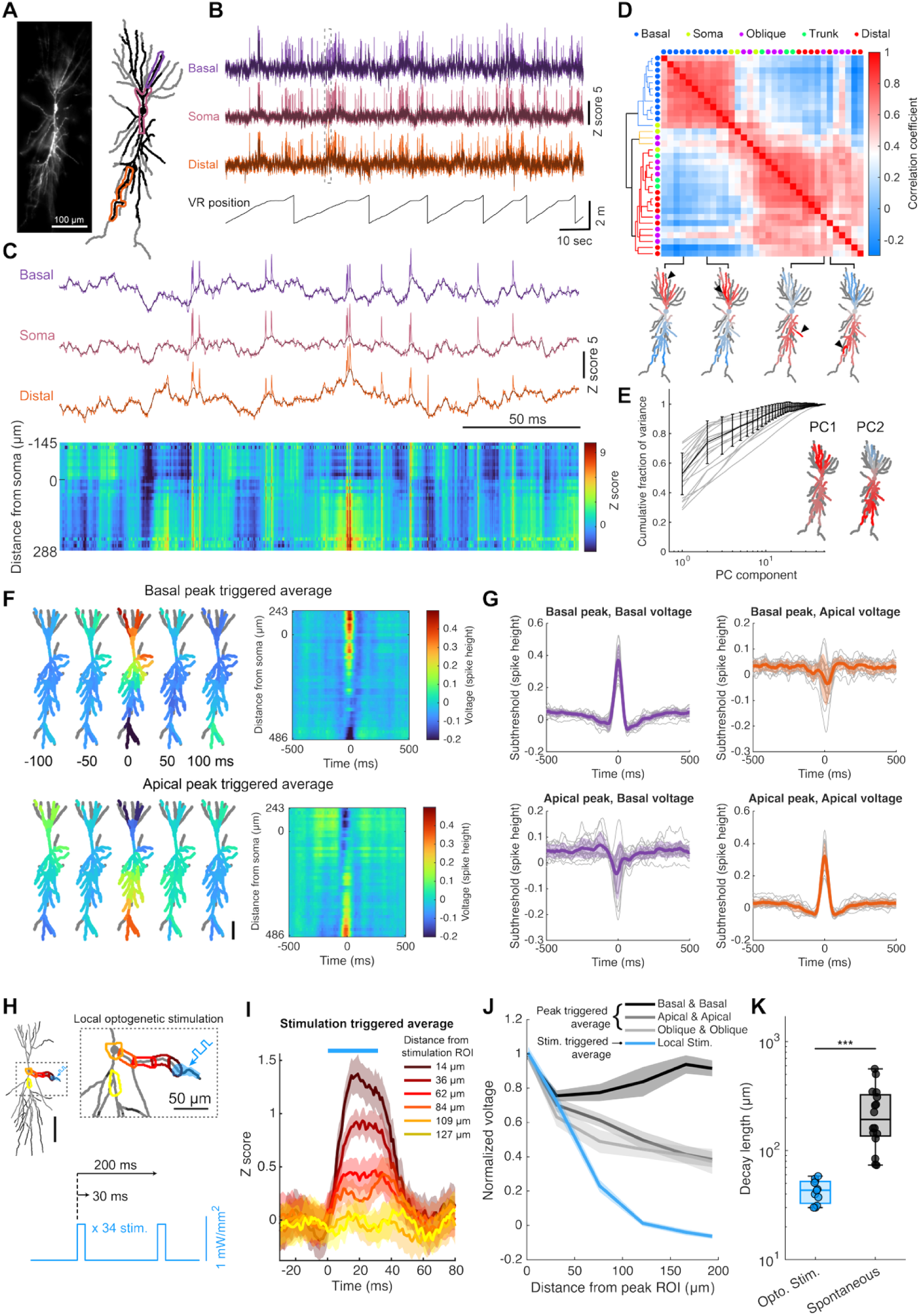
Voltage imaging reveals intra-laminar correlations of subthreshold dynamics. (A) Left: CA1 pyramidal neuron expressing Optopatch-V, imaged in vivo via DMD-targeted illumination. Right: reconstructed full dendritic structure from a z-stack (grey) and overlay of dendritic segments in the voltage-imaging field of view (black). (B) Fluorescence traces from the indicated regions in (A) and VR position (black). (C) Top: magnified view of dotted region in (B). Bottom: corresponding voltage kymograph along the basal-apical axis. (D) Correlation matrix of subthreshold dynamics among dendritic segments. Dendrite classes are indicated by colored dots. Dendritic segments are ordered by leaf order of the correlation matrix. Bottom: example correlation maps between specific trigger segments (black arrows) and the rest of the dendritic tree. (E) Cumulative fraction of variance explained by principal components (n = 20 neurons, 12 mice). The first two components explained 71 ± 14% (mean ± s.d.) of the total variance in the dendritic voltage dynamics. Inset: first two eigenvector maps showing global and see-saw components for the neuron in (A-D). Error bars indicate mean ± s.d. (F) Subthreshold peak-triggered average voltage maps, triggered by peaks in a basal dendrite (top, n = 1534 peaks) and an apical dendrite (bottom, n = 1167 peaks). Right: peak-triggered average kymographs along the basal-to-apical axis showing anti-correlation of basal and apical voltages. (G) Peak-triggered average traces for n = 20 neurons, 12 mice, with peaks detected from mice running in VR. (H) Experiment to probe responses to local dendritic stimulation in an anesthetized mouse. The stimuli comprised 30 ms pulses on a 200 ms period, repeated 34 times. (I) Example stimulation-triggered voltage responses for a single dendritic target location. Stimulus trials that evoked somatic spikes were omitted. (J) Normalized peak-triggered average voltages, comparing spontaneous peaks in basal (black), apical (grey), and oblique (light grey) dendrites (n = 20 neurons, 12 mice), and local optogenetic stimulation (blue, n = 11 experiments, 7 neurons, 5 mice). As a function of contour distance, optogenetically evoked depolarizations decayed much faster than spontaneous depolarizations. (K) Decay lengths of dendritic voltage from local optogenetic stimulation (n = 11 experiments, 7 neurons, 5 mice) and spontaneous apical subthreshold peaks (n = 16 neurons, 8 mice; Wilcoxon rank-sum test, P = 1.6 × 10^-5^). Error bars and shading in (G, I, J) indicate mean ± s.e.m. Box plots in (K) show median, 25^th^ and 75^th^ percentiles, and extrema.

We examined the correlational structure of the subthreshold dynamics. The temporal autocorrelation of the fluctuations in subthreshold voltage decayed with an exponential time-constant of 60 ± 14 ms in apical dendrites and 56 ± 19 ms in basal dendrites (mean ± s.d., *n* = 20 neurons, 12 mice, **Fig. S8**). The pairwise equal-time correlation matrix between dendritic segments showed distinct blocks corresponding to basal, soma, and apical compartments (**Fig. 2D**), and anti-correlation between basal vs. apical blocks. Principal component analysis (PCA) revealed that a global mode and a basal-apical (“see-saw”) mode together accounted for 77 ± 9% (mean ± s.d., *n* = 20 neurons, 12 mice) of the subthreshold variance (**Fig. 2E**). Both modes were consistently observed across all recorded neurons (**Fig. S9**). Thus, the subthreshold dynamics on the dendritic tree largely functioned as two or three electrical compartments.

Some theories of dendritic computation have posited that individual dendrites carry distinct subthreshold information (*26*, *27*). To test this prediction, we explored the spatial and temporal dynamics surrounding local peaks in the voltage in individual branches. In contrast to the correlational analysis which treats all time-points equally, a peak-triggered analysis captures the influence of possibly rare upward fluctuations. For each dendrite, we detected the local subthreshold voltage peaks and then constructed peak-triggered average movies of the surrounding voltage dynamics (**Fig. 2F** and **Movie S5**). Peak-triggered averages showed broadly distributed peri-peak dynamics across basal dendrites, and, separately, across distal apical dendrites. We typically observed anti-phase dynamics between apical vs. basal (in *n* = 15 of 20 neurons, **Fig. 2G**), suggesting a trans-laminar contribution to balanced excitation and inhibition (*14*, *28*).

We very rarely observed local dendritic subthreshold depolarizations (spanning ∼30 μm of dendrite, lasting 1.6–4.7 s), with distinct spatial tuning compared to the somatic spiking place field (**Fig. S10**). We only observed five such dendrite-localized subthreshold place-fields, out of a cumulative 10,000 s of recording from 20 neurons. Together, these results indicate that most neural activity in CA1 pyramidal cells does not involve branch-specific subthreshold dynamics.

We next investigated the biophysical basis of the broadly correlated subthreshold dynamics. The long-range spatial correlations could reflect either a long cable length-constant or correlated laminar synaptic inputs (or both). To distinguish these mechanisms, we measured the membrane voltage length-constant in anesthetized mice by delivering pulses of blue light targeted to small dendritic segments (**Fig. 2H**). Optogenetically evoked steady-state subthreshold depolarization decayed monotonically vs. contour distance from the stimulation site, with length-constant 43 ± 10 μm (mean ± s.d., *n* = 11 branches, 6 neurons, 5 mice; **Fig. 2I**, **Movie S6**), substantially shorter than length constants measured by patch clamp in acute slices (*29–31*).

We then repeated the analysis, triggering off spontaneous apical peaks in subthreshold voltage in the same cells. These fluctuations decayed with a length constant 240 ± 150 μm (mean ± s.d. *n* = 16 neurons, 8 mice, **Methods**), significantly longer than the length constant for optogenetically evoked depolarizations (Wilcoxon rank sum test, *P* = 1.6 × 10^-5^; **Fig. 2K**). These results establish that the broad intra-laminar correlations in subthreshold voltage are more attributable to correlated drive than to long-range electrotonic coupling. A previous study reported that, in cortex, neighboring cells had strongly correlated excitatory inputs (cross-correlation *ρ* = 0.75) (*32*); thus it is not so surprising that neighboring dendrites on the same cell would have correlated inputs.

We next sought to discern the relative contributions of excitation and inhibition to the dendritic voltage dynamics. To map excitatory inputs, we calculated the footprints of positive-going excursions from resting potential. To map inhibitory inputs, we first applied a mild constant wide-area optogenetic stimulus, to broadly depolarize the neuron. This increased the driving force for inhibitory currents, amplifying their effect on membrane voltage (*32*, *33*). Then we calculated the footprints of the spontaneous negative-going voltage excursions (**Fig. S11**, **Methods**).

Putative excitatory footprints appeared primarily on basal and apical dendrites, likely reflecting feedforward CA3 and EC inputs, respectively (*34–36*). Putative inhibitory footprints appeared primarily near the soma or in apical dendrites, potentially corresponding to PV and SST interneuron inputs, respectively (*37*, *38*). Though our assignment of these voltage fluctuations to synaptic excitation and inhibition is provisional, the footprints largely follow the known axonal projection patterns of local circuits (**Fig. S11**) and are consistent with previous reports of opposite E/I polarity between perisomatic and apical dendrites (*28*).

Taken together, these results show that dendritic subthreshold dynamics *in vivo* are highly correlated between branches within each layer, but segregated between basal and apical layers, and that the balance of excitation and inhibition arises across layers.

### bAPs are the main drivers of dendritic excitations

We next examined the spiking activity. Spikes were distinguished from subthreshold events by their fast, large-amplitude waveform (**Methods**), and were separately identified in the soma and all dendritic segments. Spikes within each dendritic segment were then linked to events in adjacent segments if the peaks occurred within 1 ms. Spikes were classified by their initiation site and whether they involved a somatic spike, following the taxonomy in **Fig. 3A**. We demarcated spikes that started at the soma (SomAPs); spikes that started in dendrites and triggered a somatic spike (dAPs); and spikes that started in dendrites but failed to evoke a somatic spike (dSpikes).

**Figure 3.**
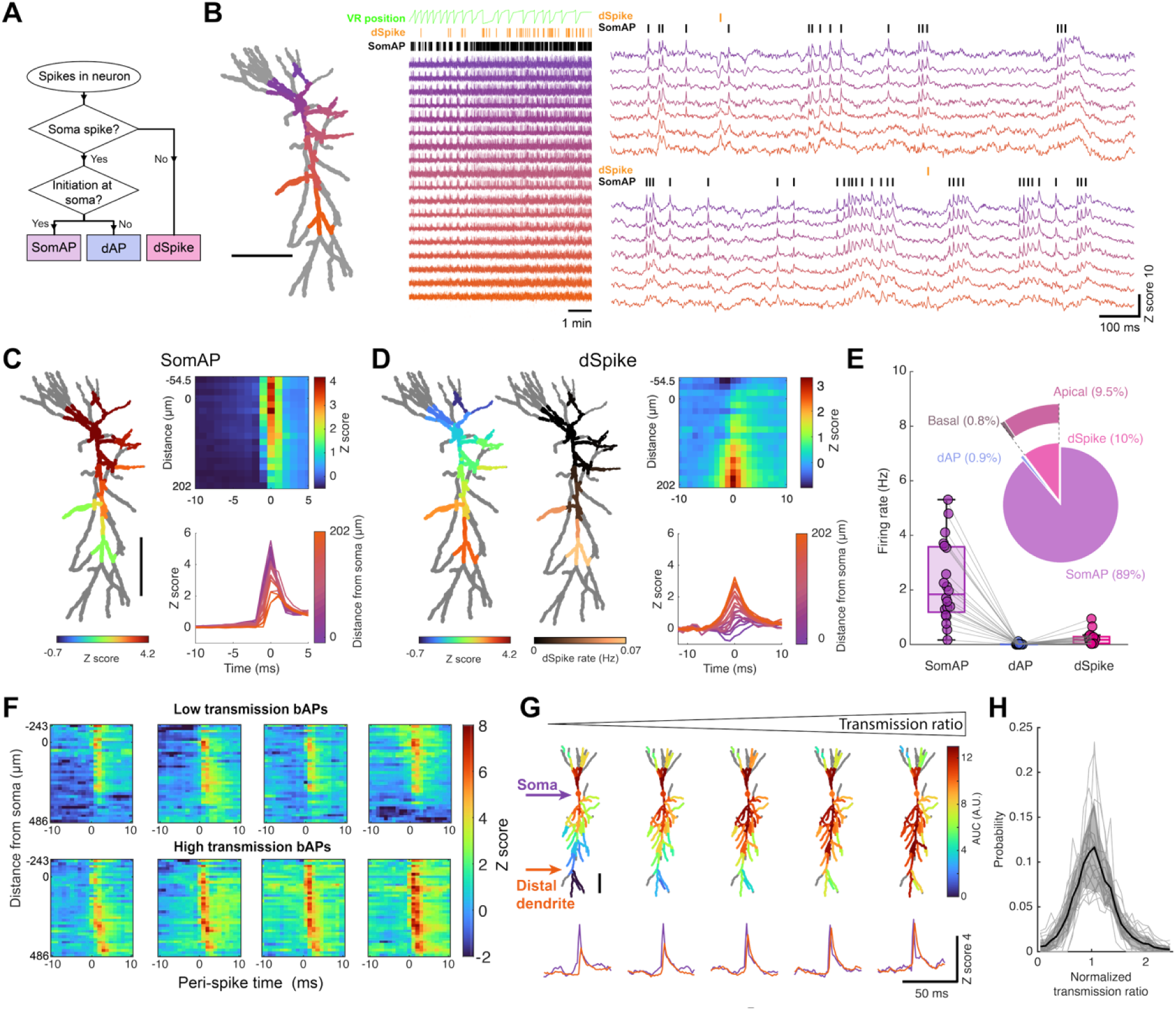
Dendritic excitations are primarily mediated by bAPs. (A) Terminology for action potentials: SomAP, soma-initiated action potential; dAP, dendrite-initiated action potential; dSpike, dendrite-localized spike not evoking a somatic spike. (B) Left: dendritic structure (grey) overlaid with dendrites in the voltage imaging field of view (colored). Middle: voltage traces from the correspondingly colored dendrites during VR navigation (location shown in green). No dAPs were detected in this neuron. Right: magnified voltage traces, showing dSpikes and SomAPs. (C) Left: SomAP-triggered average amplitude map (*n =* 658 SomAPs). Right: corresponding kymograph and voltage traces, showing spike back-propagation. (D) Left: dSpike-triggered average amplitude map (*n* = 63 dSpikes). Middle: dSpike rate map, showing dSpike prevalence in apical dendrites. Right: corresponding kymograph and voltage traces, showing coincident apical excitation and basal/somatic inhibition. (E) Rates of SomAP, dAP, and dSpike across neurons during VR navigation (*n =* 20 neurons, 12 mice). Box-whisker plots show median, 25^th^ and 75^th^ percentiles, and extrema. Inset: Prevalence of each action potential type, as a fraction of all fast spikes (somatic or dendritic). (F) Kymographs of individual bAPs showing (top): low-transmission bAPs and (bottom): high-transmission bAPs. (G) Average AUC maps (top) and voltage traces from the soma and distal apical dendrites (bottom) of bAPs sorted into quintiles by bAP transmission ratio. (H) Distribution of transmission ratios across *n* = 20 neurons, 12 mice. Shading indicates mean ± s.d.

To distinguish SomAPs and dAPs, we mapped the dendritic voltages around the time of each somatic spike. Dendritic voltage peaks clearly fell into two categories. Most events showed a distance-dependent dendritic delay relative to the soma (99% of 19,906 spikes, *n* = 20 neurons, 12 mice), corresponding to a propagation speed *V*_c_ = 230 ± 60 μm/ms away from the soma (mean ± s.d., **Fig. S6**).

In a few events (101 out of 19,906 spikes, *n* = 20 neurons, 12 mice), the dendritic voltage peaked earlier than expected from the mean propagation delay, and in some cases before the soma spiked (**Fig. S12**). These events were classified as dAPs. SNAPT analysis on a dAP-triggered average movie showed that dAPs initiated in the distal apical dendrites, propagated toward the soma, and then spread to basal and oblique dendrites (**Fig. S12D** and **Movie S7**). dAPs were rare, only 1.0 ± 0.5% of somatic spikes (mean ± s.e.m., *n =* 20 neurons, 12 mice). Out of 18 neurons where we optogenetically stimulated apical dendrites, only one showed optogenetically evoked dAPs (**Fig. S13**), though all showed optogenetically evoked SomAPs and bAPs, even with distal apical stimulation. These findings indicate that fast dendritic spikes are not a major contributor to neuronal integration.

We also observed dSpikes that did not trigger somatic spikes (10 ± 2.1% of events, mean ± s.e.m., 1,920 dSpikes out of 21,826 spike events, *n* = 20 neurons, 12 mice, **Fig. 3B**). **Figs. 3C** and **3D** compare the spatial footprints of the SomAPs and dSpikes. In dSpikes, the voltage upstroke in the apical dendrites coincided with a strong somatic hyperpolarization, indicative of somatic inhibition (**Movie S8**). We infer that this somatic inhibition prevented the soma from spiking upon arrival of the incoming dSpike. This motif resembled a greatly sped-up version of the see-saw subthreshold mode (compare **Fig. 3D** to **Fig. 2F**), suggesting that perhaps the same presynaptic cell populations were involved in both, i.e. driving strong apical excitation and perisomatic feed-forward inhibition. **Fig. 3E** shows the relative proportions of the different kinds of neuronal excitations. The greater prevalence of dSpikes vs. dAPs suggests that feed-forward somatic inhibition serves as a potent brake preventing dendritically initiated spikes from reaching the soma.

Since SomAPs were the most common events and prior studies showed that bAP propagation can trigger regenerative dendritic activity (*7*, *23*, *36*, *39–41*), we next examined bAP propagation following SomAPs. bAPs showed variable efficiency of propagation into the distal apical dendrites (**Fig. 3F**). We quantified this effect via the transmission ratio, defined as the ratio of the bAP area under the curve (AUC) in the apical dendrites (> 160 μm from the soma) to that at the soma (**Fig. 3G**, **Fig. S14**, **Methods**). The distribution of transmission ratios was unimodal (**Fig. 3H**), indicating a continuum of bAP transmission efficiencies rather than a clear demarcation into successes and failures. The mean cross-correlation of bAP AUC across distal dendrites was 0.72 ± 0.04 (mean ± s.e.m., n = 52 apical segments, 127 pairs from 20 neurons, 12 animals), suggesting broadly shared variations in bAP amplitude. We did not observe evidence of branch-specific bAP attenuation or amplification. These results show that, as for the subthreshold voltages, the dominant variation in bAP propagation was across lamina rather than between branches within a layer.

### Apical depolarization enhances bAP propagation

We asked what patterns of pre-spike subthreshold voltages facilitated or inhibited bAP propagation. We sorted spontaneous bAPs by their transmission ratio and then calculated spike-triggered averages voltage profiles of the bottom 20% and top 20% of bAPs (**Fig. 4A**). Apical hyperpolarization preceded low-transmission bAPs, and apical depolarization preceded high-transmission bAPs (*n* = 391 bAPs, Pearson correlation coefficient: 0.36, *P* = 1.21× 10^-13^, **Fig. 4B**). This correlation persisted across all bAP transmission ratios and across all recorded neurons (*n* = 20 neurons, 12 mice, **Fig. 4C**). The apical bAP transmission ratio correlated with pre-spike subthreshold depolarization in apical dendrites, but not in basal dendrites (**Fig. 4C**), and this correlation emerged ∼50 ms before the spike (**Fig. 4D**).

**Figure 4.**
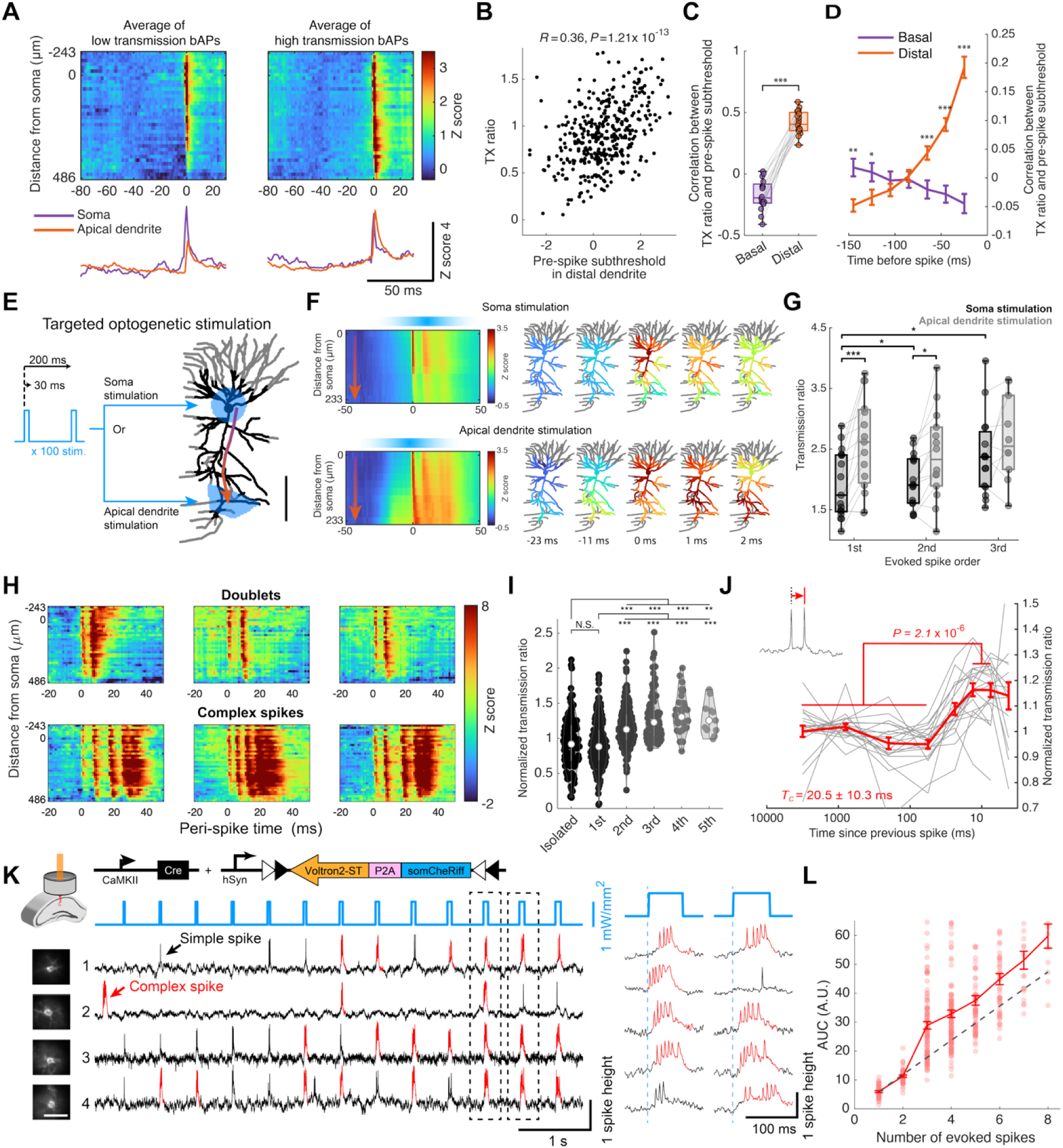
Apical voltages modulate bAP propagation. (A) Spike-triggered average kymographs and voltage traces, triggered by low-transmission bAPs (left, *n =* 22 spikes) and high-transmission bAPs (right, *n =* 22 spikes) recorded in the example neuron from Fig. 3F-G, in a mouse navigating in VR. Apical hyperpolarization preceded low-transmission bAPs and apical depolarization preceded high-transmission bAPs. (B) Transmission (TX) ratio as a function of distal apical pre-spike subthreshold voltage (mean from 10 ms to 3 ms before spike, in dendrites 349–486 μm from the soma; *n* = 391 spikes, *R*, Pearson’s correlation coefficient; *P*, two-tailed *t*-test). (C) Correlation between bAP apical transmission ratio and pre-spike subthreshold voltage in the basal and apical dendrites (*n =* 20 neurons, 12 mice; paired two-tailed *t*-test, *P* = 4.6 × 10^-10^). (D) Correlation between bAP apical transmission ratio and pre-spike subthreshold voltage as a function of pre-spike time. Apical depolarization correlated with bAP transmission within ∼50 ms before the spike (*n =* 20 neurons, 12 mice; paired two-tailed *t*-test, \**P < 0.05,* \*\**P < 0.01, \*\**\**P < 0.001*). (E) Protocol for optogenetic stimulation at either the soma or apical dendrite in an anesthetized mouse. (F) Average voltage traces triggered by the first spike after stimulation onset (soma stimulation: *n* = 95 spikes; dendrite stimulation: *n* = 73 spikes). Left: kymograph along the arrow in (E). Right: peri-spike voltage map. The pre-spike subthreshold depolarization was largest in the stimulated region. Apically evoked spikes showed higher transmission for the first bAP than somatically evoked spikes. (G) Transmission ratio for 1^st^, 2^nd^ and 3^rd^ bAPs evoked by soma (black) or apical dendrite (grey) stimulation (*n =* 17 neurons, 11 mice; paired two-tailed *t*-test, \**P <* 0.05, \*\**P <* 0.01, *** *P <*0.001). (H) Kymographs of spontaneous spike trains in an awake mouse showed that among the first 3 spikes, successive bAPs propagated further along apical dendrites. (I) Transmission ratio as a function of spike order within spontaneous spike trains (isolated and 1^st^–5^th^ spike order groups: *n* = 140, 217, 214, 84, 34, 11 spikes, respectively, Wilcoxon rank sum test, \*\**P* < 0.01, \*\*\**P* < 0.001). (J) Normalized transmission ratio plotted against interval from previous spike (decay constant *T*c = 21 ± 10 ms, mean ± s.d.; *n =* 20 neurons, 12 mice; paired two-tailed *t*-test, *P* = 2.1 × 10^-6^). For each neuron, the transmission ratio was normalized by the average transmission ratio of spikes with no preceding spikes for 300 ms. (K) CA1 neurons expressing soma-targeted Optopatch-V were imaged through a hippocampal cannula window in an anesthetized mouse, and excited with blue-light pulses of increasing duration. Left: voltages in four neurons, complex spikes shown in red. Right: expanded traces from dotted boxes. (L) Relationship between somatic voltage AUC and number of optogenetically evoked spikes. Dotted line shows linear extrapolation from the AUC of a single spike. Points show individual stimuli. Data from *n* = 1142 stimuli, 28 neurons, 3 mice. Box-whisker plots in (C) and (J) show median, 25^th^ and 75^th^ percentiles, and extrema.

To test whether pre-spike depolarization causally facilitated stronger bAPs, we optogenetically stimulated either the soma or distal dendrites in anesthetized mice (**Fig. 4E**). Voltage profiles triggered off the first evoked spike showed that distal stimulation led to more efficient bAP propagation compared to somatic stimulation (soma-stimulated transmission ratio: 1.7 ± 0.7; apical-stimulated: 2.3 ± 0.9, mean ± s.d, *n* = 17 neurons, 11 mice, paired two-tailed t-test, *P* = 4.4× 10^-4^; **Fig. 4F,G**). Together, these results demonstrate that pre-spike apical depolarization promotes more robust backpropagation of action potentials.

Studies in brain slices showed that at resting potential, apical A-type potassium (K_V_) channels shunt bAP propagation, but that apical depolarization inactivates these channels, facilitating propagation (*18*, *39*, *42*, *43*). We interpret our observations that both spontaneous and optogenetically evoked apical depolarization facilitated bAP propagation as likely evidence of this phenomenon *in vivo*.

Optogenetic stimuli of 30 ms duration—either apical or soma-targeted—typically evoked trios of spikes at ∼100 Hz (**Fig. 4F**). For soma-targeted stimulation, bAP propagation increased with spike order (**Fig. 4G**). The transmission of the third somatically evoked bAP became comparable to that of the first apically evoked bAP. The transmission of apically evoked bAPs did not increase with spike order. We observed a similar short-term facilitation of bAP propagation for spontaneous spike bursts (**Fig. 4H, I**). The window of this bAP enhancement lasted for 21 ± 10 ms after each bAP (mean ± s.d., *n* = 20 neurons, 12 mice; **Fig. 4J**). Similar short-term facilitation of bAP propagation has been observed in L2/3 neurons *in vivo* (16). Prior experiments in hippocampal slices showed that bAP-induced depolarization can transiently inactivate apical A-type K_V_ channels, clearing the way for subsequent bAPs to propagate more effectively and thereby to engage dendritic Na_V_ channels (*18*, *39*, *42*, *43*).

### bAPs trigger apical plateau potentials

Strong depolarization of apical dendrites can activate voltage-gated calcium channels and, in conjunction with glutamatergic inputs, NMDA receptors (*7*, *36*, *42*, *44*). Together these channels can drive dendritic “plateau potentials” which last hundreds of milliseconds and which drive spike bursts or complex spikes at the soma (*23*, *45*). Prior experiments in acute slices suggested that bAP-mediated dendritic sodium spikes were a necessary trigger for these plateau potentials (*7*, *18*, *23*, *36*, *41*), so we tested this hypothesis *in vivo*.

We classified spontaneous spikes into simple spikes (SSs) and complex spikes (CSs). CSs were defined as events containing three or more spikes with inter-spike interval (ISI) ≤ 20 ms and occurring on top of a large subthreshold depolarization (**Fig. S15**, **Methods**). During spontaneous CSs *in vivo*, we indeed observed a gradual buildup of apical depolarization coincident with the arrival of bAPs. This buildup typically continued until the apical voltage reached a plateau potential after 3–4 bAPs (**Fig. 4H,I** and **Fig. S16**). This observation is consistent with the idea that several bAPs in quick succession are needed to trigger the dendritic plateau. However, this correlational observation does not establish causality.

To test the causal role of bAPs in evoking apical plateau potentials, we delivered somatically targeted optogenetic stimulation pulses of varying durations to trigger different numbers of spikes in anesthetized mice (**Fig. 4K**). To enhance measurement throughput, we expressed soma-targeted Voltron2 and soma-targeted CheRiff and performed measurements through a cannula and an implanted window which exposed the dorsal surface of CA1 (*22*). Although we could not image dendritic voltage directly using this preparation, we used the evoked voltage AUC at the soma as an indicator of dendritic plateau potentials. Optical triggering of ≥ 3 somatic spikes led to a CS waveform at the soma and a corresponding jump in the somatic voltage AUC (*n =* 28 neurons, 3 mice; **Fig. 4L**). We interpret this jump as evidence of apical plateau potentials evoked by the third bAP. We then administered ketamine, an NMDAR antagonist. Ketamine abolished the optogenetically evoked CSs (**Fig. S18**), supporting the view that NMDAR currents are a necessary component of these events (*18*, *24*, *36*).

We applied the same optogenetic stimulus protocol to putative interneurons, which we classified based on soma diameter, after-spike depolarization (ADP), and peak ISI (**Fig. S19** and **Methods**) (*46*). We did not observe CSs under any stimulus durations, and the voltage AUC increased linearly with the number of evoked spikes, consistent with the absence of plateau potentials in this cell class.

In anesthetized mice with implanted prisms, distal dendritic stimulation was more effective at evoking CSs than somatic stimulation (56 ± 31% of spikes occurred within CSs for distal stimulation vs. 27 ± 13% for soma stimulation, *n =* 7 neurons, 6 mice, paired two-tailed t-test, *P* = 0.024; **Fig. S17**). Taken together, these results indicate that strong bAP propagation driven by consecutive somatic spikes or by distal pre-spike depolarization can trigger regenerative dendritic excitation, likely dependent on NMDA receptor activation, ultimately leading to CSs.

We then compared the spontaneous subthreshold voltage dynamics along the apical-basal axis leading up to CSs vs. SSs. To our surprise, we did not observe any significant differences which could predict whether the subsequent event would be a CS or an SS (**Fig. S15** and **Movie S8**). We also did not detect distinct pre-spike subthreshold dynamics of CSs that induced place-fields vs. those that did not (**Fig. S16**). The CS vs. SS spatiotemporal voltage dynamics only diverged after the first spike. Thus, the bifurcation between SS vs. CS *in vivo* may reflect aspects of the synaptic input that arrive after the first somatic spike (at which point we cannot distinguish synaptic vs. internally generated contributions to the membrane potential); or the bifurcation may depend on variables not observed in our experiments such as the fine-grained structure of synaptic inputs, E/I balance, or modulatory signals.

### Sustained spiking suppresses bAP propagation and complex spikes

Previous studies in acute slices showed that dendritic Na_V_ channels can amplify bAP propagation (*47*), and that upon sustained spiking slow Na_V_ inactivation prevented bAPs from triggering plateau potentials and CSs (*6*, *8*, *18*, *48*, *49*). To test whether slow Na_V_ inactivation affected bAP propagation *in vivo*, we delivered 500 ms blue stimuli at the soma in anesthetized mice, to induce tonic firing at 24–104 Hz (**Fig. 5A**). As in our prior experiments (**Fig. 4E, F**) the first bAP was relatively attenuated and the 2^nd–^5^th^ bAPs propagated successively more efficiently into the apical dendrites and triggered a plateau potential in the apical dendrites and a complex spike at the soma (**Fig. 5B**, **b1**; **Fig. 5C**). Later bAPs were again attenuated (**Fig. 5B**, **b2**; **Fig. 5C**). Analysis across multiple neurons revealed that bAP amplitudes transiently increased in a ∼15–60 ms window after somatic stimulation onset (**Fig. 5D**) and then decreased again, i.e. that the dendritic back-propagation acted as a spike-rate accelerometer. A similar attenuation-propagation-attenuation motif was observed in CA1 pyramidal cells in acute slices (*18*) and in L2/3 *in vivo* (*16*).

**Figure 5.**
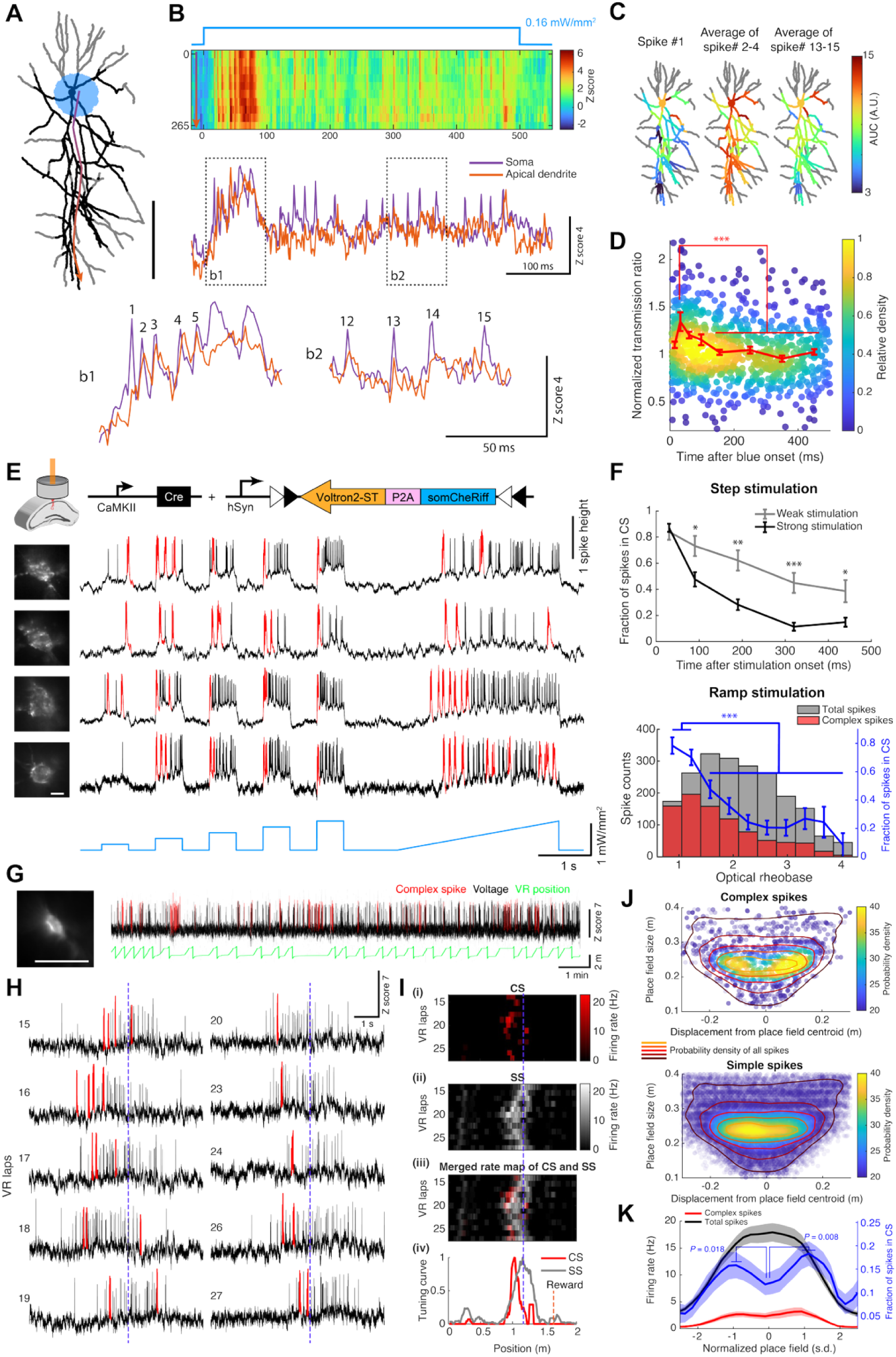
Sustained spiking reduces bAP propagation and suppresses complex spikes in the center of place fields. (A) Reconstructed neuron (grey) and dendritic segments in the voltage imaging field of view (black). A 500 ms optogenetic stimulus was applied to the soma in an anesthetized mouse. (B) Top: kymograph of voltage dynamics along the soma-apical axis indicated by the arrow in (A). Bottom: voltage traces from soma and apical dendrites. Boxed regions (b1 and b2) are expanded below, showing variation in bAP amplitude in the apical dendrite. (C) AUC maps of spike #1, the average of spikes #2-4, and the average of spikes #13-15 from (B), showing short-term facilitation and long-term suppression of bAP propagation. (D) Transmission ratio relative to stimulation onset (*n =* 927 spikes, 19 neurons, 10 mice; Wilcoxon rank sum test, *P* = 5.2 × 10^-4^). For each neuron, the transmission ratio was normalized to the average transmission ratio of attenuated spikes (spikes occurring ≥ 250 ms after stimulation onset). (E) Spiking responses from four representative CA1 pyramidal neurons expressing soma-targeted Optopatch-V in an anesthetized mouse, imaged through a cannula window. Complex spikes (CSs) are highlighted in red. (F) Top: fraction of spikes in CSs during weak (grey, < 1.5 × optical rheobase) and strong (black, > 1.5 × optical rheobase) step stimulation (n = 52 neurons, 4 mice; paired two-tailed t-test, *P < 0.05, **P < 0.01, ***P < 0.001). Bottom: total spike count (grey), CS (red), and fraction of spikes in CSs (blue) as a function of optical rheobase during ramp stimulation (*n =* 52 neurons, 4 mice; paired two-tailed t-test: weak vs. strong stimulus: *P* = 9.0× 10^-5^). (G) Example image and voltage trace of a place cell expressing soma-targeted Optopatch-V in a mouse running in VR. CSs are shown in red. (H) Enlarged voltage traces within the place field, showing CS prevalence near the edge of the place field. Blue dotted lines mark the mean place field centroid, averaged over all laps. (I) Firing-rate maps of (i) CSs (red), (ii) SSs (grey), (iii) overlay, and (iv) corresponding tuning curves for the cell shown in (G, H). (J) Top: density of CSs in place fields as a function of displacement from the place field centroid and size of the place field (*n =* 3841 spikes, 48 place fields, 39 neurons, 7 mice). Contour lines indicate probability density. Bottom: same plot for SSs (*n =* 24,118 spikes, 48 place fields, 39 neurons, 7 mice). (K) Firing rate of all spikes (black), CSs (red), and the fraction of CSs (blue) as a function of displacement from the lap-by-lap centroid normalized by place field size. Place field sizes of 20–32 cm were included in this analysis (*n =* 28 place fields, 24 neurons, 7 mice; paired two-tailed *t*-test: front vs. middle, *P* = 1.8× 10^-2^; middle vs. rear: *P* = 7.9× 10^-4^). Shading indicates s.e.m.

We next explored the impact of this modulation in bAP propagation on CS formation. We delivered soma-targeted step optogenetic stimuli (500 ms duration) at varying intensities, followed by a 3 s ramp (**Fig. 5E**). CSs were more frequent at the start of each step stimulus than later in the pulse, and the CS rate decreased more rapidly for stronger optical stimuli. During 500 ms step stimulation, weak step stimuli evoked more CSs while strong stimuli evoked more SSs (fraction of spikes in CSs during the 2^nd^ half of the weak stimulus: 0.50 ± 0.07; strong: 0.13 ± 0.03; mean ± s.e.m., *n =* 52 neurons, 4 mice, **Fig. 5F**). Similarly, the ramp stimulation evoked more CSs early in the ramp and more SSs later in the ramp (fraction of spikes in CSs during weak portion of the ramp: 0.78 ± 0.04; strong portion: 0.3 ± 0.03, mean ± s.e.m., *n =* 52 neurons, 4 mice, **Fig. 5F**). Thus, weak or transient somatic stimulation favors CSs, provided that the stimulus is long enough to trigger at least 3 somatic spikes in quick succession.

### Dendritic biophysics shape place field structure

The major excitatory input to CA1 place cells is peri-somatic feedforward input from CA3, which gradually increases toward the center of the place field (*23–25*, *50–53*). We hypothesized that the ramping somatic excitation in place fields might recapitulate the patterns of SSs and CSs we observed with optogenetic soma-targeted ramp stimuli.

To test this hypothesis, we mapped the distributions of CSs and SSs within place fields in mice expressing soma-targeted Optopatch-V (**Fig. 5G, H**). To accommodate lap-to-lap variations in firing, we measured the center and width of the place field on each lap (**Methods**) and used these to establish a normalized place-field axis. CSs preferentially occurred at the front and back of the normalized place field, compared to SSs which occurred in the middle (**Fig. 5J**). We investigated the fraction of CSs in place fields of typical size (20–32 cm). The fraction of spikes in CSs was significantly higher at the edges of the place field (paired two-tailed t-test, front vs. middle, *P* = 1.8× 10^-2^; middle vs. rear: *P* = 7.9× 10^-4^, **Fig. 5K**), consistent with previous reports showing that bursts are preferentially localized to the edge of the place field (*54*, *55*). Symmetric triangular optogenetic stimulation ramps in anesthetized mice recapitulated this pattern of CSs and SSs (**Fig. S20**). We interpret the switch from CS to SS firing in the middle of the place field as a consequence of dendritic Na_V_ inactivation under sustained high-rate bAPs. Together, these experiments show how the dendritic biophysics modulates the structure of neuronal spiking throughout a place field.

## Discussion

We investigated dendritic voltage dynamics and their influence on spike output in CA1 pyramidal cells of mice navigating in a VR environment. Dendritic subthreshold dynamics were low-dimensional, dominated by global and see-saw (basal-apical) modes. Spontaneous subthreshold fluctuations were correlated over a contour distance ∼6-fold larger than the biophysical length constant (**Fig. 2J**), indicating that these dynamics are likely driven by laminar inputs which are largely shared between branches.

Approximately 99% of somatic spikes originated near the soma, which has a lower AP threshold than the dendrites (*56*) (the axon was not visible in our measurements so we could not distinguish initiation in the soma from initiation in the axon initial segment). Apical dendritic spikes (dSpikes) occurred at ∼10% of the somatic spike rate, but these events were almost always accompanied by hyperpolarization at the soma, which blocked somatic spiking (**Fig. 3D**). While the functional role of dSpikes remains uncertain, experiments in brain slices have suggested that dSpikes may contribute to synaptic plasticity in apical dendrites (*44*, *57*, *58*). Only very rarely (∼1% of somatic spikes) did a fast dendritic excitation trigger a somatic spike. Thus, the local circuit dynamics appear structured to actively suppress somatic spike generation by fast dendritic excitations; but to permit activation of fast dendritic excitations by back-propagating action potentials.

Back-propagating action potential amplitudes also primarily varied across lamina along the apical-basal axis, rather than between dendritic branches within layers. During a high-frequency somatic spike train, bAP propagation showed short-term facilitation and steady-state suppression (i.e., a spike-rate accelerometer), likely due to fast dendritic K_V_ inactivation and slow Na_V_ inactivation, respectively (*43*, *47*). Conjunction of three or more closely-spaced bAPs with apical synaptic excitation triggered dendritic plateau potentials, which then drove complex spikes at the soma; whereas sustained somatic spiking above ∼40 Hz was not effective at triggering dendritic plateau potentials, likely due to dendritic Na_V_ inactivation. This biophysically determined window for bAP propagation led to distinct tuning curves for simple and complex spikes within place fields. The position-dependent balance of CS vs SS firing may play a role in stabilizing place fields (*25*, *59*).

Our work confirmed the *in vivo* relevance of several mechanisms which had been proposed based on brain slice experiments. These include the temporal window for bAP propagation governed by the interplay of A-type Kv and Na_V_ channels (*6*, *40*, *41*, *46*); activation of apical NMDARs by conjunction of bAPs and synaptic inputs (*18*, *36*); and driving of complex spikes by apical plateau potentials (*24*, *25*).

Our *in vivo* experiments also showed several differences from our prior voltage imaging measurements in acute brain slices (*18*). First, the distribution of bAP transmission ratios was unimodal *in vivo* (**Fig. 3H**), but bimodal – showing clear successes and failures – in slices. This difference likely reflects the highly variable availability of dendritic ion channels under *in vivo* conditions. Second, CSs could be triggered purely by somatic stimulation *in vivo*, whereas in slices they required either concurrent drive of somatic spiking and apical synaptic inputs (*18*, *36*), or presence of non-physiological ions (e.g. cesium) (*24*). This difference likely stems from tonic glutamate release that primes NMDARs *in vivo*. Third, many studies evoked dendritic spikes in brain slices using glutamate uncaging (*60*), electrical field stimulation (*57*), or dendritic current injection (*6*, *56*, *61*). Our data indicate that synaptic inputs *in vivo* are only rarely strong enough to engage these dynamics.

We did not detect a significant difference in pre-spike voltage dynamics between SSs and CSs (**Fig. S15**), nor between CSs that evoked place fields vs. those that did not (**Fig. S16**). There are several possible explanations for these null results. Perhaps the glutamatergic apical inputs that are key for CS formation come after the first bAP in a bAP train; or perhaps the key variables that govern the SS vs. CS bifurcation are not apparent in the membrane voltage, but instead reflect E/I balance, dendritic Ca^2+^ or cAMP (*62–64*) concentration, or other non-observed variables. To resolve these uncertainties, future work will need to relate dendritic voltage dynamics to patterns of inputs, as reported by reporters of glutamate (*65*, *66*), GABA (*67*), and neuromodulators; and to internal signals such as Ca^2+^ (*68*) and cAMP (*69*, *70*).

By establishing the hierarchical nature of somatically driven dendritic excitations leading to complex spikes – which are the triggers for BTSP (*23*, *24*, *71–74*) – our data point toward a plasticity-centric model of dendritic excitations in CA1 pyramidal cells. Strikingly, we observed a nonlinear transition in which 3–4 closely spaced bAPs can recruit an apical NMDA-dependent plateau potential that drives a somatic complex spike, providing a concrete *in vivo* mechanism for spike-multiplet sensitivity and a natural bridge from millisecond-timescale spiking to “instructive-event” learning rules such as behavioral time-scale synaptic plasticity (BTSP), where plateau potentials act as cell-wide teaching signals over seconds-long windows. The suppression of CSs at high spike rates via apical Na_V_ inactivation suggests a biophysically grounded mechanism for Bienenstock-Cooper-Munro (BCM)-style metaplasticity (a sliding modification threshold)(*75*), that can complement slower homeostatic mechanisms such as synaptic scaling(*76*).

Complex spikes may also differentially engage downstream circuits through short-term synaptic plasticity (*77*), and thereby enable information multiplexing without altering connectivity (*78*). Our work raises the possibility that dendritic excitations could play a role in mediating circuit-wide information processing, though this remains to be established.

## Acknowledgements

We thank Michael Goard for guidance on the prism imaging preparation. We thank Andrew Preecha and Camila Bodden for technical assistance. We thank Liam Paninski, Benjamin Antin, Utku Ferah, Chase King, Amol Pasarkar, Ahmed Abdelfattah, and Eric Salter for helpful discussions. BHL acknowledges support from National Research Foundation of Korea (NRF) grant RS-2023-00248959. XW acknowledges support from an AHA postdoctoral fellowship. SEP and LDL were supported by the Howard Hughes Medical Institute. This work was supported by a Vannevar Bush Faculty Fellowship N00014-18-1-2859, the Harvard Brain Initiative, NIH grants R01-NS126043, R01-NS133755, and R01-MH117042, and Chan Zuckerberg Initiative Dynamic Imaging Grant 2023-321177.

## Author contributions

BHL designed project, performed experiments, analyzed data, and wrote manuscript. PP developed the genetic construct and helped with the experimental design. XW performed spinning disk confocal imaging to acquire dendritic structure. JDW-C developed the optical system. JX, BHL and Y-CH mapped GFAP expression in brain slices. MX and BHL built and integrated the VR system. YQ characterized Voltron2-JF608. DGI wrote instrument control software. SEP and LL synthesized and provided JF608-HTL. AEC supervised project, wrote manuscript.

## Data availability

Data are available from the corresponding author.

## Methods

### Animals

All procedures involving animals were in accordance with the National Institutes of Health Guide for the care and use of laboratory animals and were approved by the Institutional Animal Care and Use Committee (IACUC) at Harvard University. 5–7-week-old male C57BL/6 (WT) mice were housed under a reverse 12 hr light/dark cycle. Training and experiments were conducted during the animal’s dark cycle.

### Genetic constructs

Optopatch-V, described previously (*18*), comprised a chemigenetic voltage indicator Voltron2(*19*), co-expressed with the blue-shifted channelrhodopsin CheRiff (*20*), linked via a self-cleaving P2A peptide. Voltron2 and CheRiff were each fused to membrane-targeting and the trafficking motifs to enhance dendritic expression and functional membrane localization. Specifically, TS from K_ir_2.1 was incorporated as the membrane localization sequence, LR from Lucy-Rho (*21*) was used to promote trafficking to distal dendrites, and the ER export signal ER2 (FCYENEV) was appended to improve surface expression. The final construct, hSyn::DIO-LR-Voltron2-p2a-LR-CheRiff-TS-ER2, was cloned into an adeno-associated virus (AAV) backbone in the form of a Cre-dependent DIO (double-floxed inverse open reading frame) cassette, enabling selective expression in Cre-expressing neurons. The construct was packaged using AAV serotype 2/9.

For soma-targeted Optopatch-V, we used trafficking motif from Kir2.1 for both Voltron2 and CheRiff, to restrict the expression near the soma. This construct was also designed to express in a Cre dependent manner. The final construct, hSyn::DIO-Voltron2-ST-p2a-CheRiff-ST, was cloned into an AAV backbone and packaged using AAV serotype 2/1. Both Optopatch-V and soma-targeted Optopatch-V were packaged by NeuroTools viral vector core at University of North Carolina at Chapel Hill (UNC).

### Virus injections

Virus injections were performed using a microinjection pump (World Precision Instruments, Nanoliter 2010) controlled by a microsyringe pump controller (World Precision Instruments, Micro4) and beveled glass micropipettes pulled on a Sutter P1000 puller.

Mice were subcutaneously administered dexamethasone (4.8 mg/kg), carprofen (5 mg/kg) and extended-release buprenorphine (Ethiqa XR; 3.25 mg/kg) approximately 1 h prior to surgery. Mice were anesthetized with 1.5–2.5% isoflurane (Kent Scientific SomnoSuite) and positioned in a stereotaxic frame (Stoelting) with the head secured by ear bars and upper-incisor mounting. Eyes were protected with ophthalmic ointment (Alcon, Systane) and small pieces of aluminum foil for shielding from the surgical lamp. Scalp hair was removed with depilatory cream (Nair). The scalp was then disinfected with ethanol and iodine and incised just enough to expose bregma and lambda.

Bregma and lambda were aligned in the dorsoventral axis, and the medial–lateral axis was leveled by comparing z-positions at AP −2.0 mm, ML ±2.0 mm. A 0.05 mm carbide burr was used to make holes in the skull. At the injection coordinates, virus loaded glass pipettes were lowered slowly to the target injection sites.

For microprism surgery, the mixture of AAV2/9 hSyn::DIO-LR-Voltron2-p2a-LR-CheRiff-TS-ER2 (final concentration: 1.6×10^12^ GC/mL; Addgene #230988), AAV2/1 hSyn::DIO-Voltron2-ST-p2a-CheRiff-ST (final concentration: 4.8×10^11^ GC/mL), and AAV2/9 pENN.AAV.CamKII 0.4.Cre.SV40 (final concentration: 0.3–3×10^7^ GC/mL; Addgene, #105558) was injected at 3 sites in left hippocampal CA1 (AP: 1.4, 2.1, 2.8 mm; ML 1.7mm; DV: 1.0 –1.4 mm). The small portion of soma-targeted Optopatch-V was included to improve the peri-somatic voltage signal. At each site, 160 nL of virus was delivered at 60 nL/min.

For hippocampal window surgery, the mixture of AAV2/1 hSyn::DIO-Voltron2-ST-p2a-CheRiff-ST (final concentration: 8×10^12^ GC/mL) and AAV2/9 CamKII 0.4::Cre.SV40 (final concentration: 0.6–1.2×10^9^ GC/mL; Addgene, 105558) was injected at 5 sites in left hippocampal CA1. Injections were centered at AP -2.1 mm and ML 1.7 mm, with four additional sites located 0.7 mm radially from the center. Virus was delivered at DV 1.0–1.4 mm, 160 nL per site, at 60 nL/min.

Following injections, the scalp was sutured (2–3 stitches) and sealed with cyanoacrylate tissue adhesive (3M, Vetbond). Animals were monitored daily, and carprofen (5 mg/kg) was administered once per day for 3 days.

### Microprism and hippocampal window implantation

Microprism implantation was performed on mice one week after AAV virus injection. The implantation protocol was adapted from (*17*). All surgical procedures were performed under aseptic conditions.

For implantation surgery, preoperative procedure was the same as virus injection. The scalp was resected to expose a wide region of dorsal skull, including bregma and lambda. The periosteum was completely removed using fine forceps, micro curettes, or diluted hydrogen peroxide.

Microprisms (V2HPC design from (*17*); 2.5 mm height × 1.5 mm width right angle prism, with the hypotenuse coated in aluminum, Figure S1A; Tower Optical) were bonded to a 3 mm diameter coverslip using UV optical adhesive (Norland, NOA81). The prism was positioned such that its 2.5 mm × 1.5 mm face was aligned with a 0.5 mm offset from the center of the coverslip. Assemblies were UV-cured for ∼10 min and inspected for complete polymerization.

To implant the prism into the same site as the virus injection, bregma and lambda were carefully leveled as described for virus injection procedure. A small rectangular hole was made (0.5 × 1.5 mm) in the skull to introduce a 1.5 mm 45° single-edge diamond knife (Fisher Scientific, 501930985). Using the diamond knife, a vertical incision was made from the skull surface to a depth of 1.5–2.0 mm, with the longer side of the knife facing posteriorly. To minimize bleeding, the knife was slowly lowered (1 mm/min), while ensuring the cortex was not dragged along with the blade.

A 3 mm craniotomy was made after the incision using a 3 mm diameter biopsy punch and a carbide burr. The dura within the craniotomy was removed. Prism assemblies were held on a blunt 14-gauge cannula connected to vacuum suction. The prism edge was aligned to the cortical incision and slowly lowered (1 mm/min) until the coverslip contacted the skull surface. To prevent the prism from dragging the cortex, the brain was kept moist with saline, and the prism imaging plane was monitored through the periscope during implantation. If blood obscured the imaging plane, the prism was withdrawn, the site irrigated with saline, and reinsertion attempted. The prism position was then finely adjusted until the hippocampal subfields (CA1, CA3, DG) were visible through the prism. The prism was then secured to the skull using cyanoacrylate adhesive (3M, Vetbond). After curing, a stainless steel headplate (as described in (*79*)) was positioned parallel to the coverslip and bonded with dental cement (Parkell, C&B metabond, 242-3200) that was colored black with graphite.

For hippocampal window implantation surgery, the implantation protocol was adapted from (*22*). Hippocampal window implantation was performed on the same day with AAV injection. Stainless steel cannulas (3.0 mm outer diameter, 2.8 mm inner diameter, 1.7 mm height) were bonded to a 3 mm diameter coverslip using UV optical adhesive (Norland, NOA81). A 3 mm craniotomy was made using a 3 mm diameter biopsy punch and a carbide burr. The dura was removed and the cortex was then carefully aspirated using vacuum suction fitted with gel-loading tips (Genesee Scientific, 14-100). The cortex was aspirated until the corpus callosum was visible. The cannula window was then inserted into the craniotomy and lowered onto CA1 until the window touched the tissue. The cannula was then secured to the skull using cyanoacrylate adhesive (3M, Vetbond) and headplate was attached in the same way as described in the microprism implantation surgery.

After surgery, mice were placed in a warmed recovery cage (37 °C) with hydrogel supplementation. Animals were singly housed to prevent disruption of the headplate. Carprofen (5 mg/kg) was administered daily for 3 days, and animals were monitored closely throughout the postoperative period. Mice were allowed to recover for at least 7 days before behavioral training or experimental procedures.

### Immunohistochemistry for fixed brain tissue

Mice were deeply anesthetized with an intraperitoneal injection of a ketamine/xylazine mixture (ketamine, 100 mg/kg; xylazine, 15 mg/kg) for induction, and transcardially perfused with ice-cold phosphate-buffered saline (PBS), followed by 4% paraformaldehyde (PFA) in PBS (Thermo Fisher Scientific, J19943K2). Brains were removed and post-fixed in the same fixative at 4 °C for 12–24 h, then washed thoroughly in PBS. Fixed brains were sectioned at a thickness of 150 µm using a vibratome.

Free-floating brain slices were permeabilized in PBST (1% Triton X-100 in PBS) for 24 h at room temperature on a shaker, followed by blocking in 5% bovine serum albumin (BSA) in PBST for 1 h at room temperature with gentle agitation.

For GFAP immunostaining, slices were incubated with a rabbit anti-GFAP primary antibody (1:500 dilution in 1% BSA in PBST; Abcam, cat. no. 7260) for 24 h at room temperature on a shaker. Slices were then washed three times in PBST for 20 min each and incubated with a goat anti-rabbit IgG (H+L) cross-adsorbed secondary antibody conjugated to Alexa Fluor 488 (1:500 dilution in 1% BSA in PBST; Thermo Fisher Scientific, cat. no. A11008) for 2 h at room temperature with gentle agitation.

Following secondary antibody incubation, slices were washed twice for 20 min and once for 24 h in PBST at room temperature. Slices were subsequently mounted (Vectorlabs, VECTASHIELD PLUS Antifade Mounting Medium, H-1900-10) and imaged using a Zeiss Axio Scan.Z1 wide-field epifluorescence microscope.

### Microscope system

The optical system was the same as described in (*16*) (Figure 1C). Briefly, the microscope was equipped with a 488 nm laser (Coherent, OBIS, 60 mW) for optogenetic stimulation and a 607 nm laser (Opto Engine LLC, MLL-FN-607, 300 mW) for voltage imaging. The two lasers were combined using a dichroic mirror (IDEX, FF506-Di03-25x36) and passed through an acoustic-optic tunable filter (AOTF, Gooch and Housego, TF525-250-6-3-GH18A) for intensity modulation. The beam from each wavelength was then separated using a dichroic mirror (IDEX, FF506-Di03-25x36) and relayed onto one of two digital micromirror devices (DMD, Vialux, V-7000) for patterned illumination. The patterned 488 nm and 607 nm light from each DMD was then merged using a dichroic mirror, re-imaged through excitation tube lens (Thorlabs, LA1417-A) and objective lens.

The excitation and emission paths were separated by using a multiband dichroic mirror (Chroma, ZT488/607rpc-25x36). Emission fluorescence was collected by the same objective lens and passed through an emission filter (Semrock, FF01-709/167-25), and relayed by a tube lens. Two objective–tube lens combinations were used: (i) a 25x objective (Olympus, XLPLN25XSVMP2) with a 50 mm tube lens (Zeiss, Planar T* 50mm f/1.4 ZE), and (ii) a 10x objective (Olympus, XLPLN10XSVMP) with a 100 mm tube lens (Thorlabs, TTL100-A). Fluorescence images were acquired by a scientific CMOS camera (Hamamatsu, ORCA-Fusion). The camera’s clock signal (hSync) was routed to a data acquisition system (National Instruments, NI-PCIe-6343), which also generated trigger pulses for frame acquisition. The same data acquisition system generated the experiment waveforms controlling the AOTFs, DMDs, shutters, and VR. We used home-built MATLAB/C++ based software for bidirectional microscopy, Luminos (*80*) (http://www.luminosmicroscopy.com), to generate experimental waveforms, control the DMDs and acquire images.

### VR system

The virtual reality (VR) apparatus design was adapted from the Christopher Harvey lab (https://github.com/HarveyLab/mouseVR). ViRMEn software (*81*) (https://pni.princeton.edu/pni-software-tools/virmen) was used to design and emulate VR environment. Briefly, the VR environment generated from ViRMEn was projected to a parabolic screen (210 x 580 mm) which covered ∼ 215° of mouse’s visual field using a video projector (Texas instrument, DLP3010EVM-LC). Animals ran on a cylindrical treadmill (150 x 60 mm, diameter x width) that was 3d printed with polylactic acid (PLA). The virtual environment was updated at 30–60 Hz according to the treadmill rotation, measured by an optical rotary encoder (Broadcom, HEDS-5500).

A USB data acquisition device (National Instruments, NI USB-6001) was used to synchronize with microscope DAQ, read the position encoder and lick signals, and control the solenoid valve and DMD trigger. The VR DAQ triggered the blue-light DMD when the animal entered specific VR locations for position-specific optogenetic stimulation (Figure S7). To synchronize voltage imaging and VR logging, the microscope DAQ sent the frame clock to the VR DAQ. The digital output signal from microscope DAQ and behavior data such as treadmill rotation, reward, and lick detection were recorded to a computer at every update of the VR environment.

The water reward was dispensed through a blunt-tip 12G needle connected to a solenoid valve (Lee Company, LFMX0524000A) and gravity-fed from an elevated reservoir, with the valve controlled by the VR DAQ and calibrated to deliver ∼5 μL per reward. The licking behavior was detected by measuring resistance changes in an RC circuit formed between a wire soldered to the lick spout and the grounded metal headplate.

### VR training

Prior to VR training, mice were water restricted for 5 days, to a body weight decrease to 85–90%. During this period, mice received 1–1.5 mL water per day and were habituated to the experimenter. Mice were then habituated to head restraint and running on the treadmill while exposed to the training environment for 2 weeks.

Animals were head-fixed and placed ∼20 cm from the screen. The animal’s headplate was leveled ∼3 cm from the treadmill top and the animal’s body center was placed ∼11 cm posterior to the front treadmill roller.

The training environment consisted of a 2 m-long 1D linear track with a black-and-white horizontally striped floor, a vertically striped ceiling, and a repeated arrow-pattern wall to provide optic flow. A torus and cone-shaped VR objects were positioned at the midpoint of the track. A 7.5 cm black tunnel was placed at both the beginning and end of the track to provide continuity, with mice teleported from the end to the start in the middle of the tunnel. A reward cue (green-star pattern) was presented randomly at one of three locations (30, 60, or 90% of the track length), and ∼5 μL of water was delivered at each reward position. Mice that ran more than 50 m within a 30 min session were considered trained and ready for the experiment. Typically, training took 10–14 days.

During voltage imaging experiments, mice were exposed to the training environment for the first two laps and were then switched to a novel environment on the third lap. The novel environment contained the same tunnel but was more visually enriched than the training environment, with distinct pattern combinations across each third of the track and a reward cue that differed from that used during training (Figure 1C).

### Image acquisition

Voltage imaging sessions were performed at least 8 weeks after surgery. The chemigenetic voltage indicator, Voltron2, requires a fluorogenic HaloTag dye to become fluorescent. At least 12 hours before imaging, we delivered the JF608 HaloTag ligand (120 μL, 0.5 mM), prepared as described in (*19*). Briefly, 50 nmol of JF608 was dissolved in 10 μL DMSO, 10 μL Pluronic F-127 (20% w/v in DMSO), and 100 μL PBS. Mice were anesthetized with 1.5–2.5% isoflurane, and 120 μL of the dye solution was injected into the retro-orbital sinus using a 31G syringe (MedSupply, BD-324909).

Neurons located 50–300 μm from the surface of the prism or imaging window were recorded. After positioning the target neuron at the center of the camera’s field of view, a reference image was acquired. Background subtraction and a Frangi filter (https://www.mathworks.com/matlabcentral/fileexchange/24409-hessian-based-frangi-vesselness-filter) were applied to the reference image, followed by skeletonization. The resulting skeleton was then dilated using a 5–8 μm radius sphere kernel and the resulting binary pattern was sent to the orange DMD for targeted illumination (Figure S4). For optogenetic stimulation experiments, ROIs were manually drawn and sent to the blue DMD for targeted stimulation. The imaging laser (607 nm) was delivered at 100–130 mW/mm², and the stimulation laser (488 nm) at 0.1–1.2 mW/mm² at the sample plane. All voltage recordings were acquired at 1 kHz using Luminos software (*80*).

### Statistics

Statistical significance was assessed using two-tailed paired t-tests or Wilcoxon rank-sum tests as appropriate, with the tests specified in the figure legends and significance indicated as \**P*< 0.05, \*\**P* < 0.01, and \*\*\**P* < 0.001.

## Data analysis

### Structure z-stack imaging

The morphology of neurons (Figure S1B and Figure S2) was reconstructed from three-dimensional z-stack images acquired with a 0.5 μm z-step using either a spinning disk confocal or epi-fluorescence microscope. Background signal was removed using the rolling ball background subtraction function in ImageJ (*82*).

For Figure S1B, z-stack images from three different FOVs were tiled using 3-dimensional stitching algorithm implemented in Fiji (https://github.com/fiji/Stitching) and maximum projected along the z-axis.

### Dendrite tracing

The background-subtracted z-stack images were then imported to Simple Neurite Tracer (SNT) (*83*), which provides semi-automated tracing for z-stack images. Trace results such as dendrite coordinates, branch order, and radius were manually proof-read using SNT’s path editing features, saved in SWC format, and visualized in MATLAB.

### Voltage signal extraction

All image analysis was performed using MATLAB. Imaging processing procedures are illustrated in Figure S4. Starting with the raw voltage imaging movies, we first corrected the time delay artifact from rolling-shutter of the sCMOS camera. Each row of the image has a 4.88 μs delay relative to the subsequent row, potentially distorting measurements of action potential propagation velocity. We corrected this offset by digital interpolation. We next corrected transverse (xy) motion using NoRMCorre (*84*).

To mask blood vessels, we computed a lag-1 autocovariance image, 𝐶(𝑥, 𝑦) = 〈Δ𝐹(𝑥, 𝑦, 𝑡)Δ𝐹(𝑥, 𝑦, 𝑡 + 𝑑𝑡)〉_𝑡𝑡_ , where *dt* is the inter-frame interval and 〈⋅〉_𝑡_ indicates an average over time. The lag-1 autocovariance image suppresses shot-noise and highlights pixels with high temporal variance. Pixels corresponding to blood vessels were identifiable in this image by their characteristic dot-shaped footprints, and these pixels were masked for all subsequent analyses.

We then manually drew ROIs for each dendritic segment (approximately 20 × 5 μm). To extract voltage-related pixels, we performed principal component analysis (PCA) within each dendritic-segment ROI, yielding eigenvectors, eigenvalues, and corresponding fluorescence time courses (Figure S4). Eigenvectors associated with voltage-related components were manually selected and retained for signal extraction. In most cases, the first principal component represented the spatial footprint of the Voltron2 signal and exhibited fluorescence time courses correspond to neuronal voltage dynamics (i.e., spikes and subthreshold fluctuations). This procedure largely eliminated residual contributions from tissue motion, blood-flow, heartbeat, and breathing.

Occasional particulates in the immersion water produced Airy-disk-like artifact patterns in the voltage movies. To detect frames affected by these artifacts, we first defined an Airy-disk template from a representative artifact image and convolved it with the ΔF images. Pixels with correlation values greater than 0.25 were classified as candidate particle locations. Candidate particles located within 6 μm across adjacent frames were linked into trajectories using the *track* function in MATLAB. Particles detected in at least 70 frames and traveling more than 58 μm were classified as particulate artifacts. These trajectories typically showed linear, unidirectional motion. To account for potential undetected trajectory segments, we extended the particle trajectory by linear extrapolation, adding 20% to both the beginning and end. Finally, all pixels within 35 μm of the particle’s center position were excluded from subsequent analysis.

In some recordings, we observed that dendrites moved outside the patterned illumination or drifted out of the focal plane when the mice engaged in intense movement (e.g. grooming). To detect such frames, we computed the Pearson correlation between each frame of the voltage movie and a reference image defined as the average of the first 1,000 frames. Frames showing decrease in correlation to < 0.9 were excluded from further analysis.

Voltage traces were then extracted by computing the dot product between the voltage movie and the spatial footprint of each ROI. We next filtered out the contribution from photobleaching and brain motion from the trace. The photobleaching rate of Voltron2 was estimated by an exponential fit to the mean fluorescence intensity during silent periods, (i.e., periods without spiking or optogenetic stimulation).

Brain motion can shift the signal source relative to the targeted illumination and introduce small axial-focus deviations, which are not corrected by rigid-body translation. Residual motion and photobleaching components were removed by linear regression of the following signals against the fluorescence trace from each dendritic ROI: fitted bleaching trace, motion traces 𝑥(𝑡), 𝑦(𝑡), 𝑥^2^(𝑡), 𝑦^2^(𝑡), and 𝑥(𝑡)𝑦(𝑡), calculated from NoRMCorre. We scaled these components by their regression coefficients and subtracted the resulting regressors from trace for each ROI.

The contribution from VR display flicker was removed from the traces using a band-stop filter (241.7–242 Hz).

To correct for the gradual decay in signal amplitude due to photobleaching, we first coarsely detected spikes and then fit an exponential function to the spike amplitudes vs. time. The voltage traces were divided by this exponential fit.

### Spike detection

Voltage traces were high-pass filtered by subtracting a moving-median sliding window (50 frames). Spikes were then detected using the *findpeaks* function in MATLAB. The *findpeaks* algorithm identified events that exceeded a user-defined threshold and were separated by at least 3 frames from neighboring peaks. The detection threshold was manually adjusted for each recording and typically ranged from 3 to 6 z-scores.

Dendritic spike events were classified based on their temporal relationship to the nearest somatic spike (Figure 3A). A dendritic spike was classified as a dSpike if no somatic spike occurred within ± 4 ms of the event. Within dSpike events, dendritic spikes were almost always observed across different dendrites with temporal offsets of 1–2 ms. The dSpike initiation was defined as the ROI with the earliest dendritic spike in the event.

If a somatic spike occurred within 4 ms of the dendritic spike, the event was further evaluated using the soma-to-dendrite distance and the spike timing difference. To estimate the sub-millisecond timing of the spike peak, spline interpolation was used. When the dendritic spike preceded the somatic spike and satisfied the distance criterion, the event was classified as a dAP. The distance threshold was determined by the expected back-propagation delay, computed as:

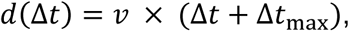

Where 𝑣𝑣 is conduction velocity (230 μm/ms, measured as described below) and Δ𝑡𝑡_max_ is one frame (1 ms). The events other than dAP were classified as somAPs. All detected somAP, dAP, and dSpike events were manually inspected to ensure accurate classification.

### Subthreshold voltage trace

To extract subthreshold voltage traces, peri-spike frames (−3 ms to +3 ms relative to each spike, including somAP, dAP, and dSpike) were removed from the voltage traces, and the missing segment was linearly interpolated using the data points at ±4 ms. The resulting trace was then smoothed using a 20 ms sliding average.

### Complex spike detection

For complex spike detection (Figure S15), both the somatic spike trace and the subthreshold trace were analyzed. Transient subthreshold bumps that exceeded the detection threshold were first identified. The detection threshold varied across neurons, typically ranging from a z-score of 3.5 to 6, corresponding to 50–60% of the spike amplitude. For each bump, we defined its duration as the interval between upward and downward crossings of a threshold z-score of 1.5. We then measured the spike ISI and spike count within this interval. If the bump contained more than 3 spikes, with the mean of ISI of the first three spikes below 20 ms, the spike train event was classified as a complex spike. Spike trains (ISI ≤ 20 ms) that occurred without accompanying subthreshold bump were classified as burst spikes (Figure S15B). All events tagged as complex spikes were manually inspected to ensure accurate classification.

### Voltage signal normalization

To compensate for nonuniform background signals and heterogeneous voltage-indicator density along dendrites, the voltage signal at each dendritic segment was normalized by the standard deviation of its subthreshold dynamics (**Fig. S5, Movie S4**). Variance arising from motion, blood vessels, and shot noise was excluded by identifying their characteristic fluorescence footprints. Using a singular value decomposition (SVD) of the voltage trace (ROIs × time), the voltage at dendritic segment *n* and time *t* of can be written as,

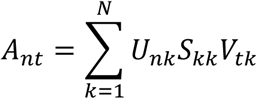

Where 𝑈_𝑛k_ denotes the spatial component, 𝑆_kk_ component weights, 𝑉_𝑡k_ the temporal components, and *N* is the number of PCs (= number of ROIs). To estimate the variance associated with voltage-related components, we examined the spatial singular vectors and selected a subset 𝐾 corresponding to voltage signals. The subthreshold variance of dendritic segment 𝑛 was then computed as,

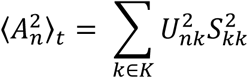

where ⟨⋅⟩_𝑡_ indicates averaging over time and 𝐾 denotes the selected voltage-related components. The voltage traces were normalized by the square root of this variance (= standard deviation, F_std_), and all dendritic voltage data presented in z-score units.

For voltage signal normalization in the movies (Movie S3–S8), the same method was used to estimate the normalization image (F_std_ image). To generate this image, the ΔF movie (pixels × time), after correction for rolling shutter, motion, and photobleaching, was used in place of the voltage trace.

For figures showing multiple neurons expressing the soma-targeted voltage indicator (Figure 4K–L, Figure S19, Figure 5E–F, Figure S20), the voltage traces were normalized by their spike height.

### Conduction velocity measurement

To measure the conduction velocity of the bAP (Figure S6E), we first computed the spike triggered average voltage at each dendritic ROI. The resulting bAP waveforms were then spline-interpolated at a 100 kHz sampling resolution to determine the sub-frame timing of the peak. Conduction velocity was measured by performing a linear regression between peak delay and distance within 200 μm apical from the soma.

### SNAPT movie

Pixelwise spike delay estimation was performed using the Sub-Nyquist Action Potential Timing (SNAPT) technique as described previously (*20*). First, spike-triggered average (STA) movies were computed. For each pixel’s time trace, the spike waveform was fitted with a quadratic spline interpolation. The sub-frame delay map was then generated by calculating the time at which the waveform reached 50% of its maximum. The delay map was converted into a spatiotemporal reconstruction of spike propagation movie by convolving it with a Gaussian temporal kernel (𝜎 = 0.05 ms). The propagation movie was then scaled by the spike amplitude map, obtained by dividing the maximum intensity projection of STA movie by the F_std_ image. The resulting propagation movie (grayscale) was mapped to a color scale (hot colormap) and overlaid on the grayscale structural image.

### Cross-correlation calculation

In Figure 2D, the subthreshold correlation between dendritic segments was calculated using a shot-noise–robust method, similar to that described in (*16*). Briefly, because the shot noise is statistically independent across neighboring pixels, the cross-correlation between odd-pixel and even-pixel signals provides an estimate of the underlying voltage correlation that is uncontaminated by shot noise. To implement this, the footprint of each dendritic ROI was divided into odd and even pixels in a checkerboard pattern. The divided patterns were then multiplied (via dot product) with the voltage movie to extract the corresponding voltage traces. The cross-correlation of subthreshold was calculated only during silent periods, i.e., intervals in which neither spikes nor dendritic plateau potentials occurred. The cross-correlation between subthreshold of dendritic segment ROIs 𝑖𝑖 and 𝑗 was calculated as following,

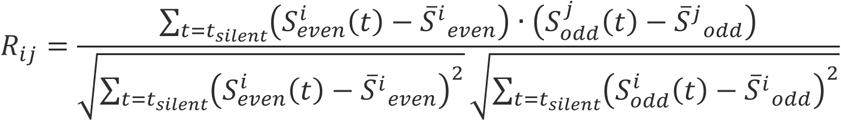

Where 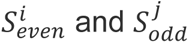 correspond to the subthreshold traces of the even pixels of ROI 𝑖 and odd pixels of ROI 𝑗 , respectively. The shot-noise-robust approach affected the autocorrelation values (*i* = *j*) but not the cross-correlation values (*i* ≠ *j*).

### Power spectrum analysis

Subthreshold signals from each compartment class (e.g., basal, soma, apical, and distal) were first averaged. The subthreshold power spectrum was calculated only during silent periods. Because the subthreshold traces contained discontinuities due to spikes, dendritic plateau potentials, and artifact-rejected frames, the power spectrum was computed using a moving-window Lomb–Scargle periodogram (3000 frame window with 1500 frame overlap), which accommodates missing samples (Figure S9). The resulting power spectrum was then normalized by the variance of the subthreshold dynamics. Theta power was quantified as the mean normalized power in the 5–10 Hz frequency band.

### Subthreshold peak detection

The subthreshold peaks were detected using the *findpeaks* function in MATLAB. Peaks that exceeded 40% of the spike height in the subthreshold trace of each dendritic segment during silent periods were classified as subthreshold peaks. When multiple subthreshold peaks occurred within 100 ms, only the largest peak was used.

For the calculation of peak-triggered average decay with distance (Figure 2K), the subthreshold peaks detected at apical dendrites were averaged and the peak-triggered average voltage amplitude was fitted with an exponential function as a function of the contour distance from the triggering location. Neurons whose maximum inter-dendritic distance was shorter than 220 μm were omitted from the analysis.

### Local subthreshold event detection

Local subthreshold events were detected from subthreshold voltage traces extracted from dendritic ROIs (Fig. S10). The global and see-saw modes calculated by PCA were first regressed out from the subthreshold traces. Candidate local subthreshold events were identified as bump-like voltage deflections whose amplitudes exceeded a z-score of 1.5 and that were detected concurrently in at least two neighboring ROIs. All detected events were manually inspected in the voltage movie to exclude artifacts (e.g., from blood flow), and only validated events were retained for analysis.

### Mapping dendritic footprints of subthreshold excitation and inhibition

Dendritic footprints of subthreshold inhibition were measured using weak wide-field optogenetic depolarization (0.03 – 0.1 mW/mm^2^). Wide-field blue light illumination was applied periodically (2 s on, 2 s off) to depolarize neurons. Because the reversal potential of inhibition is approximately -75 mV (*32*), depolarization increased the driving force for inhibitory currents, thereby enhancing the voltage response upon synaptic inhibition. Excitatory events were identified during the blue-off peak period as positive deviations exceeding a z-score of 0.7 from baseline. Inhibitory events were identified during the blue-on period as negative deviations falling below a z-score of −0.7 relative to the median voltage during stimulation.

To generate voltage footprints, subthreshold voltage movies were constructed by excluding frames near spikes, linearly interpolating the omitted values, applying a moving-average (20 frames), and normalizing by the F_std_ image described above. Voltage footprints were then computed as spatial profiles of voltage changes associated with putative excitatory or inhibitory events.

### Template-based filtering

In Movie S3 and Movie S6, to filter blood vessel and background noise, we reconstructed the movies using voltage footprint templates. The templates were derived from the dominant eigenvectors from subthreshold movies and the spike-propagation movies. Because the templates from different sources were not orthonormal, we performed Gram–Schmidt process to orthogonalize them. The reconstructed movie was computed as 𝑆𝑆^𝑇^𝑀, where 𝑆 is a template matrix with dimensions (number of pixels) × (number of templates), and 𝑀𝑀 is the vectorized movie with dimensions (number of pixels) × (number of frames). The reconstructed movie accounted for ∼75% of the variance of the original movie.

### Transmission ratio measurement

The transmission of the bAPs was measured by the ratio of dendrite voltage area under the curve (AUC) divided by the soma voltage AUC (Figure S14). The AUC was the area between the voltage trace and a local baseline. The baseline was estimated from the silent period (≥ 20 ms away from any spiking events), linearly interpolated across silent segments and smoothed with a 500 ms moving average. Because spikes in dendrite were wider than soma, somatic AUC was integrated from -1 to 1 ms relative to the spike, and dendritic AUC was integrated from -2 to 3 ms relative to the average time delay from the somatic spike. For calculation of transmission ratio, AUC of apical dendrites farther than 160 μm from the soma was averaged and divided by the somatic AUC.

In Figure 3H and Figure 4J, the transmission ratio was normalized to the average transmission ratio of spikes that were not preceded by any spike within the preceding 300 ms.

In Figure 5D, the transmission ratio was normalized to the average transmission ratio of spikes evoked ≥ 250 ms after stimulation onset.

### Variance measurement

In Figure 2E and Figure S8, for measurement of variance accounted for by each principal component, the eigenvalue of each PC was divided by sum of all eigenvalues.

### Firing map and tuning curve

To calculate the firing maps in Figure S7, Figure S16 and Figure 5I, the 2 m VR track was divided into 150 spatial bins (1.3 cm per bin). The periods during which the mice were running at speeds greater than 4 cm/s were analyzed. We calculated the mean firing rate for each bin and each lap. For the complex spike firing map (Figure 5I), individual spikes within complex spikes were counted separately. Firing maps were then smoothed using a 5-bin sliding average with circular boundary conditions. The tuning curve was then calculated by averaging the firing maps across laps.

### Spatial information

The spatial information (SI) quantifies how reliably a neuron fires at specific locations (Figure S7) and was calculated as described in (*85*). SI was computed as follows,

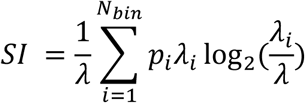

Where 𝜆 is the mean firing rate, 𝜆_𝑖_ is the mean firing rate at bin 𝑖, 𝑝_𝑖_ is the probability that the mouse is in bin 𝑖, and 𝑁_𝑏𝑖𝑛_ is the total number of bins (150 bins).

### Circular permutation test for position tuning

In Figure S10E, position tuning of local dendritic subthreshold voltage was assessed by computing the mean voltage in spatial bins along the VR track. Statistical significance was evaluated using a circular permutation test (100,000 shuffles) in which the voltage trace was circularly shifted by a random offset independently within each lap, preserving lap structure and temporal autocorrelation while disrupting the relationship between voltage and position. For each spatial bin, p-values were computed as the fraction of shuffled values exceeding the experimental value.

### Place field detection and size measurement

For detection of place fields, we required that (i) the SI of a neuron was significantly higher than the random shuffling, and (ii) the place field firing rate exceeded a defined threshold. The significance of SI was assessed by applying a random circular permutation to the bin index within each lap of the firing-rate map. A neuron was considered a potential place cell if its SI exceeded the 95^th^ percentile of 1,000 sets of random permutations. For neurons classified as potential place cells, place fields were identified as regions in which the mean firing rate exceeded 1.5 Hz and the peak firing rate exceeded five times the mean firing rate of the lowest 30% of bins. The preliminary range of place fields was defined as spatial bins with firing rates greater than two times the mean firing rate of the lowest 30% of bins. In Figure 5J,K, more detailed place field properties were computed for each lap that the mouse traversed the place field.

The centroid of the place field on lap 𝑙 was calculated as follows,

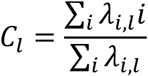

The size of place field on lap 𝑙 was calculated as follows,

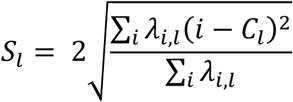

Where 𝜆_𝑖,𝑙_ denotes the firing rate at bin 𝑖 on lap 𝑙. All detected place fields were manually inspected.

### Neuronal type classification

The AAV2/9 CamKII 0.4::Cre.SV40 virus expressed Cre primarily in CA1 pyramidal neurons (PYRs) but also exhibited leaky expression in inhibitory interneurons (INTs). Because the inhibitory neurons have distinct spike waveforms and morphological characteristics, we classified them using after depolarization (ADP), ISI profiles and the size of the soma (Figure S19). The ADP for each neuron was calculated from the spike-triggered average voltage, measured as the mean voltage 4-6 ms after the spike. The peak ISI was defined as the maximum of the ISI histogram. K-means clustering along these three parameters (k = 2) was used to separate putative PYRs and INTs. The ADP of INTs was substantially smaller than that of PYRs (Figure S19A). PYR exhibited peak ISIs of ∼ 8 ms, whereas INTs showed peak ISIs of ∼ 15 ms. The soma diameter of PYRs was approximately 15 μm, whereas that of INTs was approximately 30 μm.

### Spike AUC calculation under soma stimulation

In Figure 4K-L and Figure S19, the AUC was computed by integrating the voltage signal from −9 ms to +10 ms relative to the spikes evoked by optogenetic stimulation.

### Optical rheobase measurement

In Figure 5E,F, the optical rheobase was measured by averaging the blue-light intensity required to evoke the first two spikes during somatic ramp stimulation. Stimulation intensities exceeding 1.5 × rheobase were classified as strong stimulation.

## Supplementary movie captions

**Movie S1. 3D rendering of the mouse brain showing an implanted microprism and axonal projections from CA3 and EC to CA1.** A 1.5 × 1.5 × 2.5 mm prism was implanted to the hippocampal CA1 to provide side-on view of CA1 pyramidal neurons (red). The three-dimensional morphologies of CA3 neuron axon (orange, AA0999, DOI:10.25378/janelia.7821965) and an EC neuron axon (green, AA1047, DOI:10.25378/janelia.7822316) were overlaid to illustrate their axonal projections to CA1. The microprism was positioned parallel to axonal pathways to minimize damage. Related to Figure 1.

**Movie S2. Z-stack imaging through a microprism in a live mouse and dendritic tracing.** Neurons expressing Voltron2-JF608 were imaged using a spinning disk confocal microscope. Z-stack images were acquired at 0.5 μm steps. Depth from the prism surface is indicated in the top left. The reconstructed dendrite tracing (yellow) is overlaid onto the image. Scale bars (red, green, blue): 100 μm. Related to Figure 1.

**Movie S3. ΔF/F_std_ voltage movie during a mouse navigating a virtual environment.** Top: a filtered, artifact-corrected ΔF/F_std_ voltage movie was mapped to a color scale and overlaid on the grayscale structure image; depolarization is shown in red, hyperpolarization in blue and spikes in yellow. Middle: overhead view of the virtual reality showing the mouse’s real-time position. Bottom left: mouse’s point of view. Bottom right: real-time somatic voltage trace. Related to Figure 1.

**Movie S4. SNAPT movie.** A Sub-Nyquist Action Potential Timing (SNAPT) movie showing the bAP propagation. The movie was reconstructed from the spike-triggered average movie of spikes evoked by optogenetic stimulation at the soma (blue contour). The algorithm is described in Methods. Related to Figure 1G.

**Movie S5. Peak triggered average movie.** A subthreshold peak-triggered average movie, triggering off the basal dendrite peaks (top) and apical dendrite peaks (bottom). Red arrows indicate the dendritic branches used for triggering. Peaks at basal dendrite were accompanied by the troughs at apical dendrite, and *vice versa*. Related to Figure 2F, G.

**Movie S6. Local stimulation triggered average movie.** A stimulation-triggered average voltage movie showing the subthreshold voltage profile evoked by local dendritic stimulation. Thirty-four repetitions of a 30 ms optogenetic stimulation delivered to the dendritic tip (blue region). The stimulus-triggered average -ΔF/F_std_ voltage movie was mapped to a color scale and overlaid on the grayscale structure image. Related to Figure 2H, I.

**Movie S7. dAP SNAPT movie.** A SNAPT movie showing the propagation of a dendrite-initiated spike. The movie was reconstructed from the dAP-triggered average movie (31 dAPs averaged). Related to Figure 3.

**Movie S8. SS, CS, and dSpike triggered average movie.** SS, CS, and dSpike triggered average movies and traces of three neurons. The ΔF/F_std_ movies were mapped on a color scale (shown in the color bar) and overlaid on the grayscale structure image. The movies show similar pre-spike subthreshold dynamics preceding SS and CS. The dSpike-triggered average movie shows the somatic hyperpolarization coincident with dSpikes at apical dendrites. Related to Figure 3, Figure 4, and Figure S15.

## Supplementary figures

**Figure S1.**
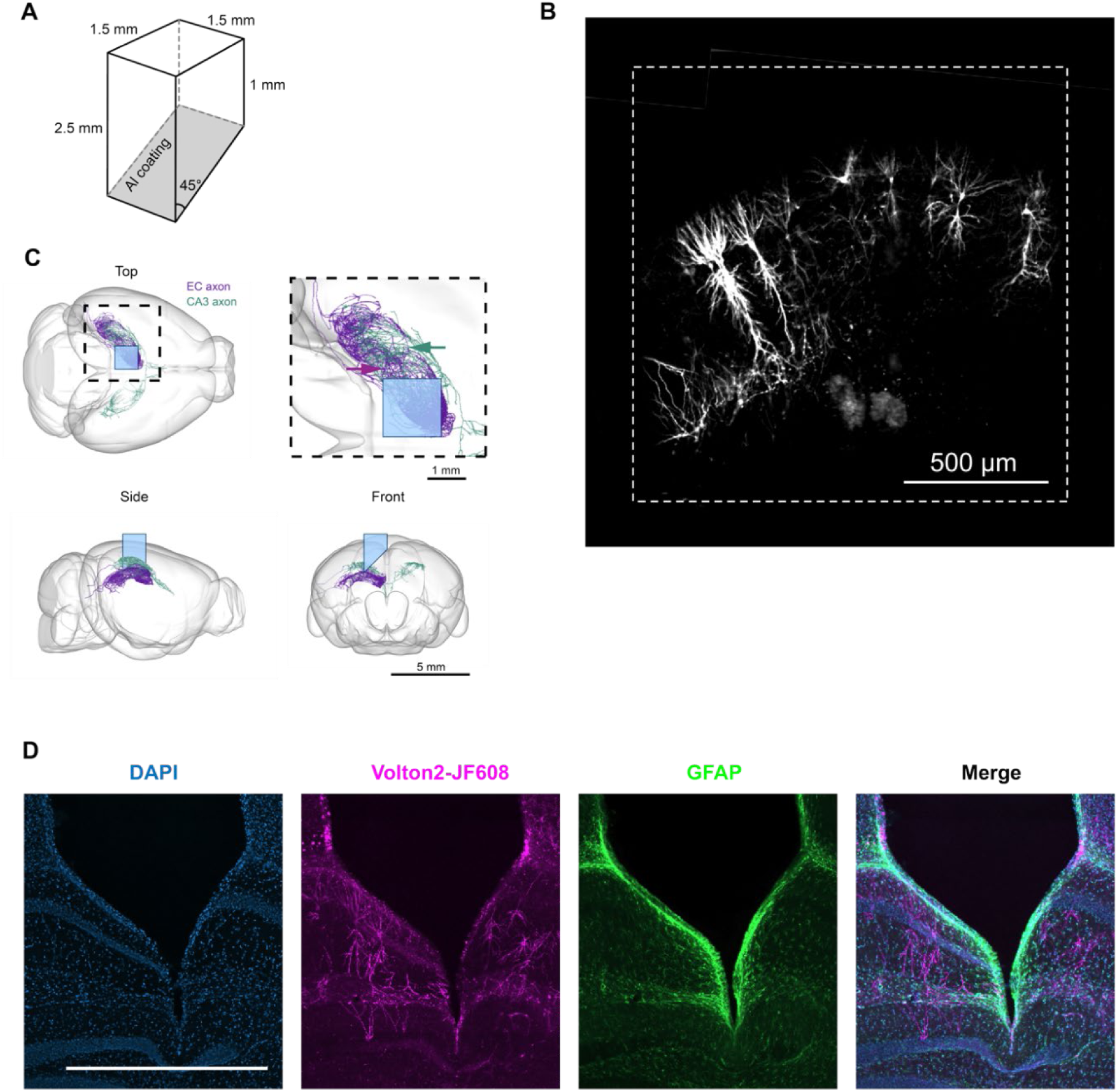
Microprism implantation. **(A)** Dimensions of the microprism. **(B)** Image of hippocampal CA1 through a microprism in a live mouse brain. Spinning disk confocal z-stacks of Voltron2-JF608 fluorescence were laterally tiled. The image shows a maximum-intensity projection. The dotted line indicates the boundary of the prism. **(C)** Schematic illustration of axon projections from CA3 (cyan) and entorhinal cortex (EC, magenta) to CA1. The microprism was inserted parallel to the laminar projection (arrows) of CA3 and EC axons. Morphological data were imported from the MouseLight Neuron Browser (https://ml-neuronbrowser.janelia.org/; EC neuron: AA1047, DOI:10.25378/janelia.7822316; CA3 neuron: AA0999, DOI:10.25378/janelia.7821965). **(D)** Images of a histological section stained for DAPI (blue), Voltron2 (magenta), and astrocyte processes (GFAP, green). Scale bar 1 mm.

**Figure S2.**
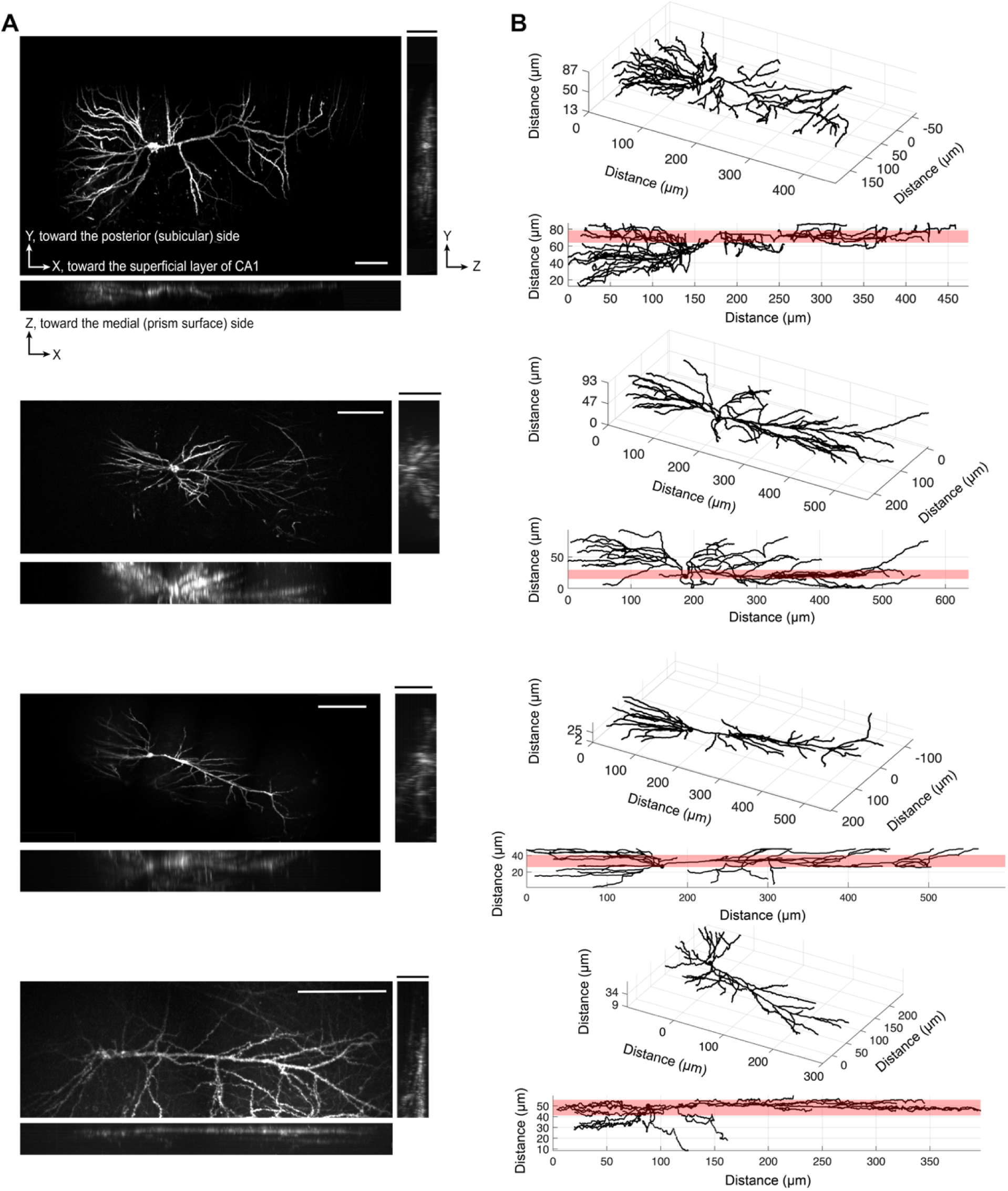
Examples of neuronal morphology. **(A)** After the voltage recording, z-stack images were acquired with a spinning disk microscope. The z-stacks were then maximum-intensity projected along the x-, y-, and z-axes. Scale bars, 100 μm. **(B)** Neurite tracings of the neuron in (A). The full width at half maximum of the voltage-imaging point spread function, centered on the focal plane, is highlighted in red.

**Figure S3.**
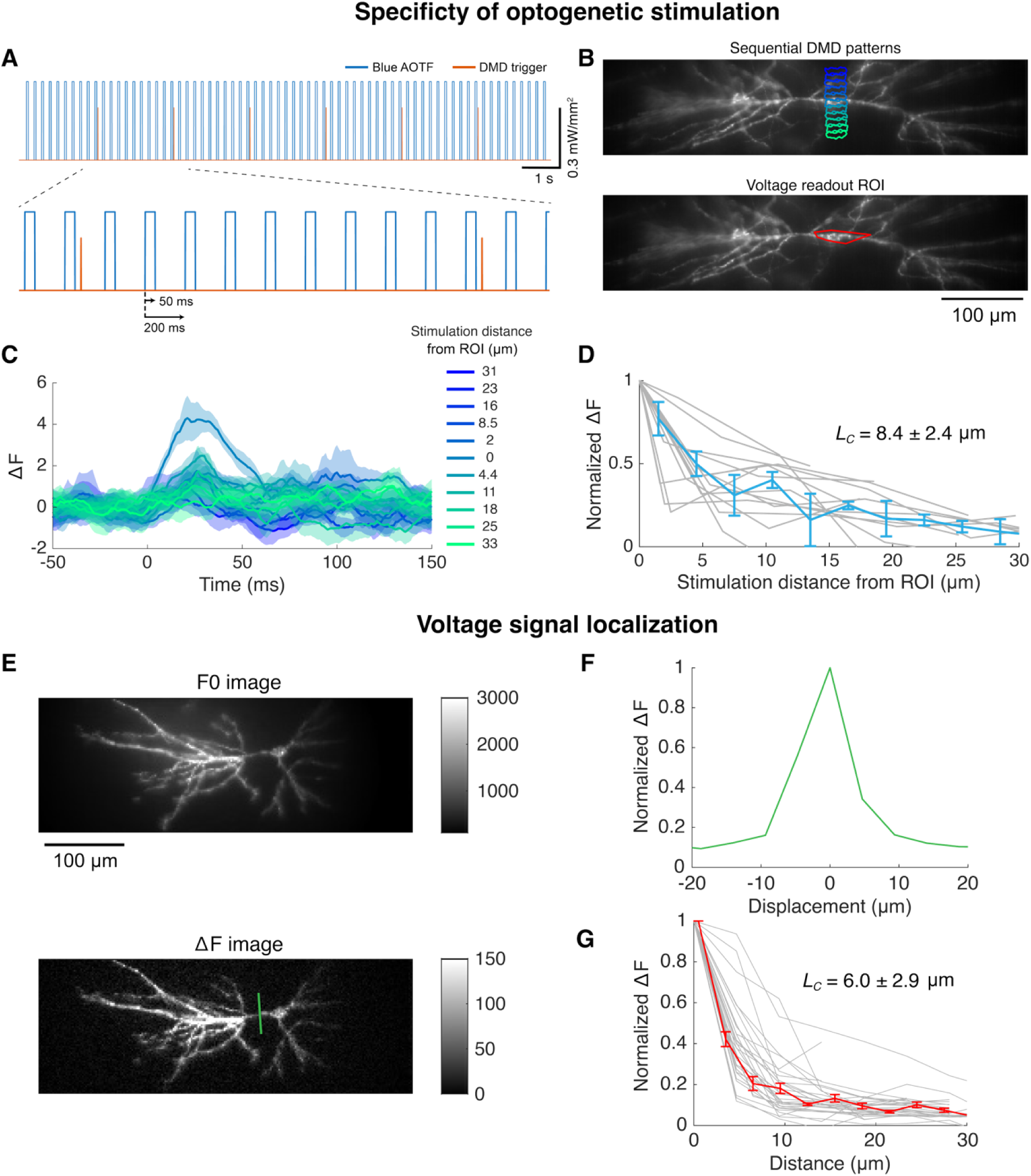
Blue -light specificity and voltage signal localization. **(A)** Experimental protocol for measuring optogenetic stimulation specificity. Stimulation patterns were preset on the DMD, and a digital input triggered the DMD to advance to the next pattern. For each pattern, 10 repetitions of 50 ms step stimulation were delivered at 5 Hz. **(B)** Examples of DMD patterns (top, blue-to-green ROIs) and the readout ROI (bottom, red). **(C)** Average voltage responses to the stimulation patterns shown in (B). **(D)** Average amplitude within 10–30 ms after stimulation onset, plotted as a function of stimulation distance (*n =* 6 branches, 3 neurons, 2 mice). Two lines are plotted for each stimulus location, corresponding to two transverse displacement directions. **(E)** Top: Mean fluorescence image. Bottom: Average spike amplitude image. **(F)** Fluorescence deviation profile along the green line in (E). **(G)** Voltage decay as a function of distance from the dendrite (*n =* 15 neurons, 7 mice). Two lines are plotted for each dendrite, corresponding to two transverse displacement directions.

**Figure S4.**
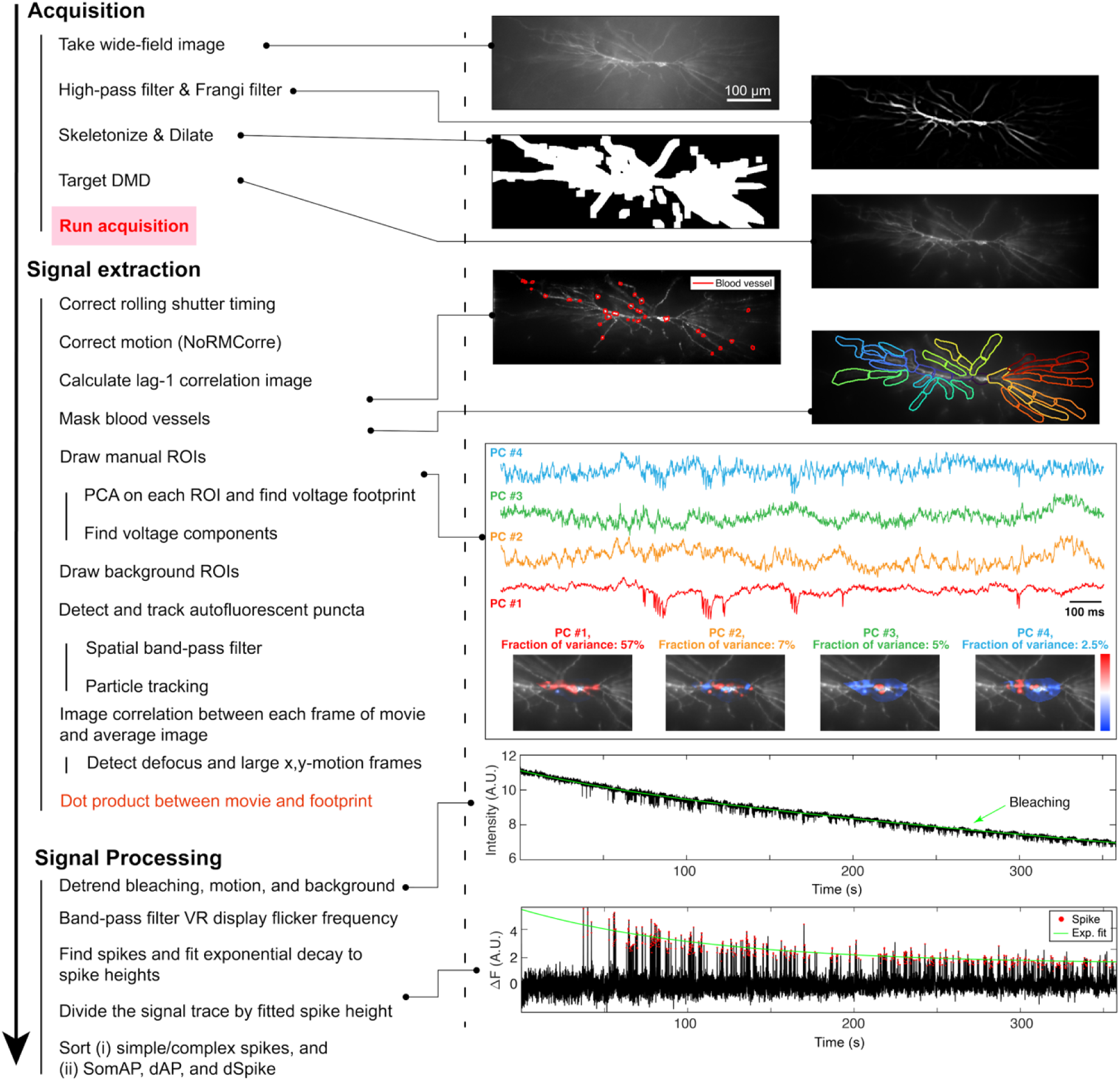
Image processing pipeline. First, a wide-field reference image was acquired. The reference image was high-pass filtered in space and processed with a Frangi filter. The resulting skeleton image was binarized and dilated and was then projected by the DMD to target the illumination to the neuron. After acquiring the voltage movie, we corrected for row-dependent time shifts caused by the camera’s rolling-shutter. We corrected for sample motion by NoRMCorre. We constructed an image that highlighted fluctuating sources (avoiding contamination from shot-noise) via the temporal lag-1 covariance of the time-stack (Methods). Pixels containing blood vessels were identified and masked. To extract the voltage signal, ROIs (20–30 μm in length) were manually drawn along the dendrites. PCA analysis within each ROI identified the footprint of the voltage signal. The temporal trace and spatial footprint of principal components (PCs) are shown in the PCA analysis panel, with eigenvectors displayed in color overlaid on the grayscale F_0_ image. The PCs corresponding to voltage signals were manually selected. Background ROIs corresponding to out-of-focus neurons were drawn separately, and their signals were later regressed out of the voltage trace. Frames contaminated by particulates in the immersion water or by large out-of-plane motion were omitted from analysis. The voltage fluorescence traces were calculated by taking the dot product of voltage footprints with the voltage movie. Bleaching, motion, and background signals were regressed out from the voltage traces, and artifacts from the VR monitor were removed using a band-stop filter (241.7–242 Hz). To correct for the gradual decay in spike heights due to photobleaching, each trace was divided by an exponential fit to the spike amplitudes. Finally, spike events were detected in each ROI and classified as simple or complex spikes. Each spike was further classified as (ii) SomAP, dAP, or dSpike.

**Figure S5.**
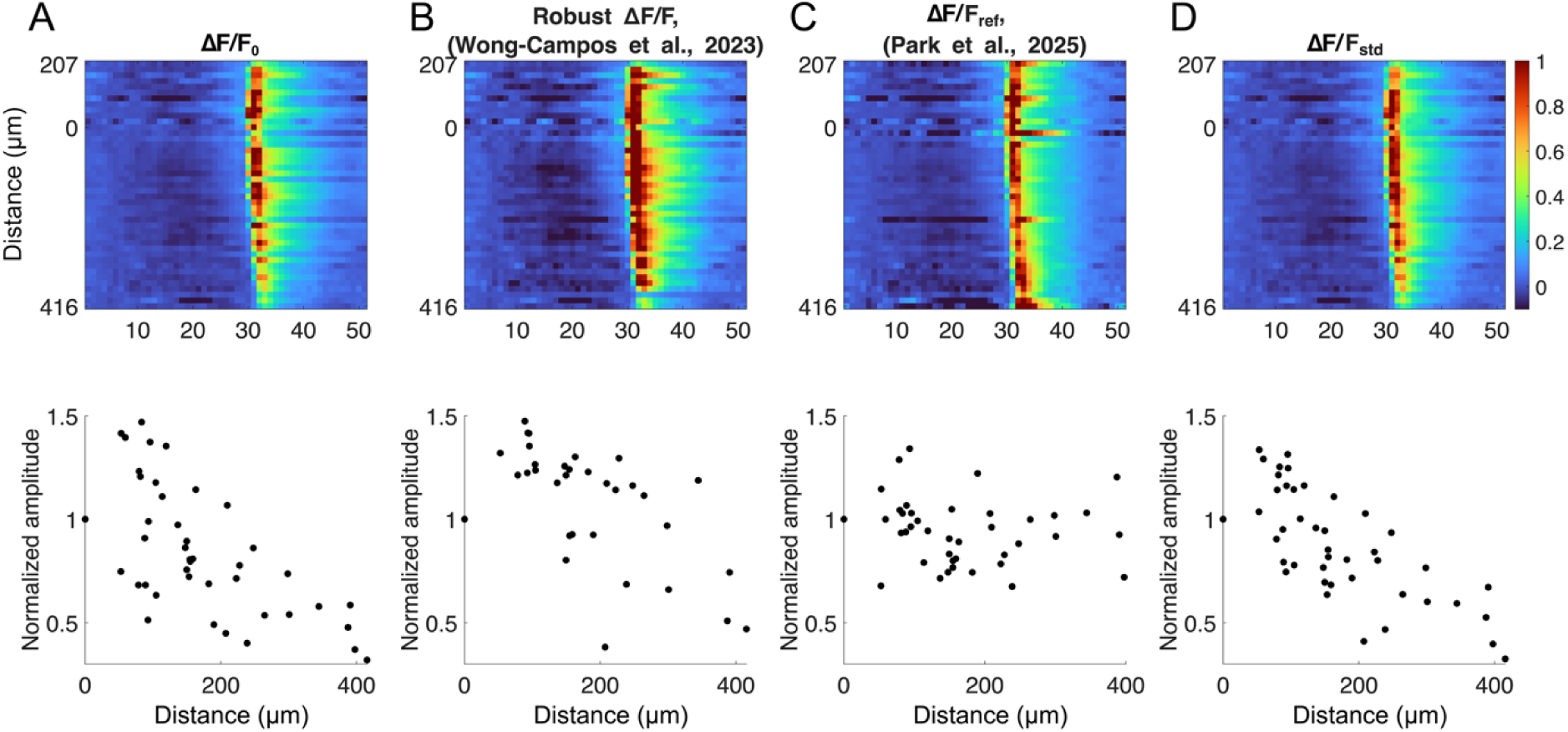
Scaling of voltage signals across dendritic branches. A challenge in dendritic voltage imaging is to compare the amplitudes of voltage signals from different subcellular locations, which might have different GEVI expression levels and fluorescent backgrounds. Kymographs of a spike-triggered average bAP were computed with four normalization methods. **(A)** Normalization by mean fluorescence intensity (F_0_). This is the widely used ΔF/F_0_ scaling. **(B)** Normalization by robust ΔF/F (*16*). The upper hull of a pixel-wise scatter plot of ΔF vs. F_0_ is fit with a line, whose slope determines the local voltage sensitivity. **(C)** Normalization by after-spike depolarization, F_ref_ (*18*). **(D)** Normalization by an estimate of the subthreshold standard deviation, that is insensitive to contamination by shot noise (F_std_; **Methods**). Normalization by standard deviation was used whenever data were reported as a z-score (**Methods**). Bottom: bAP amplitude decay vs. apical distance from the soma, measured with the different normalization methods. Method (D) showed the smoothest decay and had the lowest scatter.

**Figure S6.**
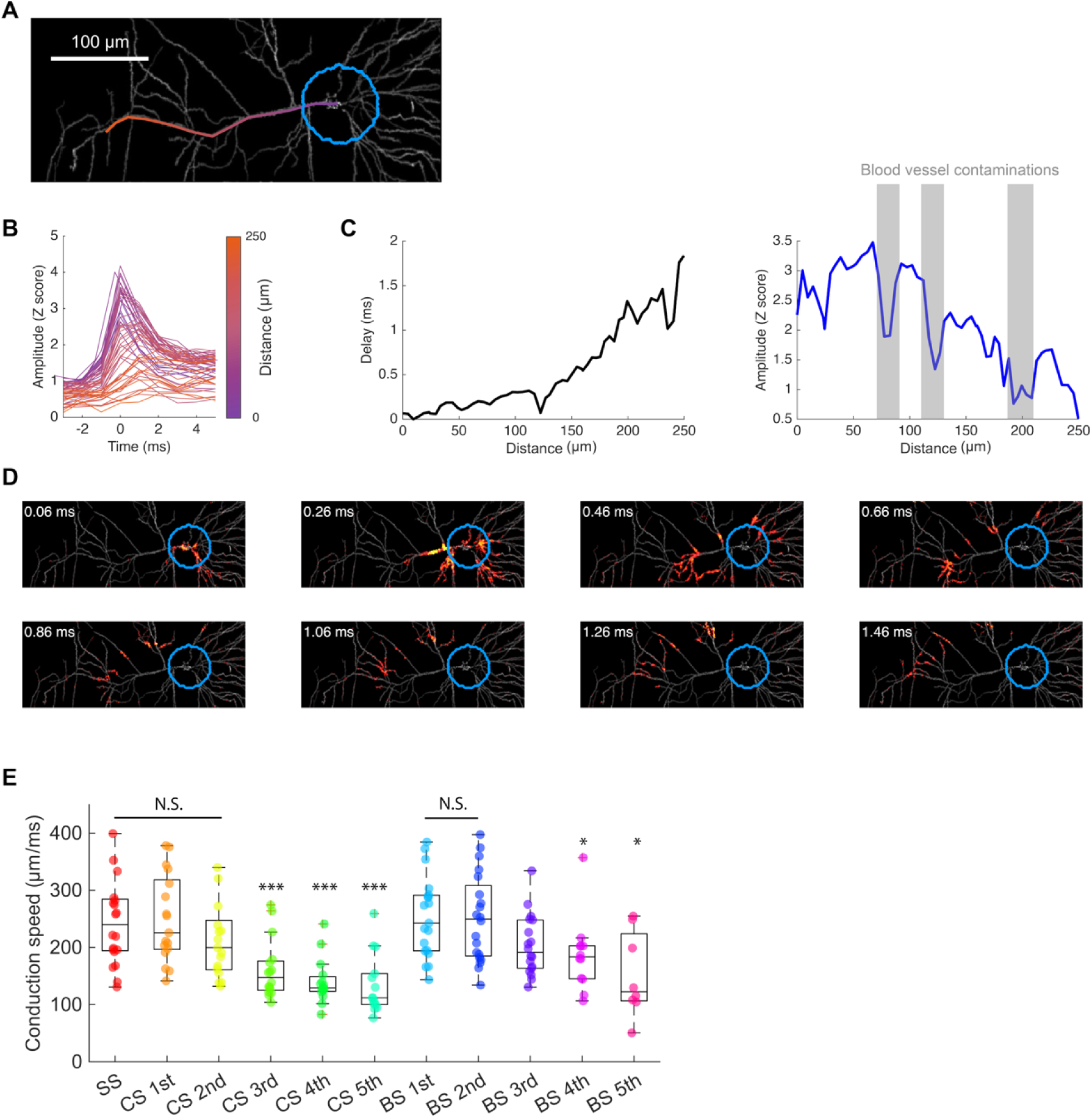
Speed of back-propagating action potentials. **(A)** 30 ms blue light stimulus pulses were delivered to the soma and proximal dendrites (blue circle). Scale bar, 100 mm. **(B)** Spike-triggered average voltage traces from the soma to apical dendrites along the line shown in (A). **(C)** Left: Spline-interpolated peak delay times along the line in (A). Right: Spike amplitudes along the line in (A). Dips reflect contamination from blood vessels (grey bars). **(D)** Snapshots from a SNAPT movie (Movie S4) showing the propagation of the bAP. **(E)** Conduction speed of simple spikes (SS), individual somatically generated spikes within complex spikes (CS), and burst spikes (BS). Stars indicate significance relative to SS (n = 20 neurons, 12 mice, paired two-tailed t-test, *P < 0.05, ***P < 0.001).

**Figure S7.**
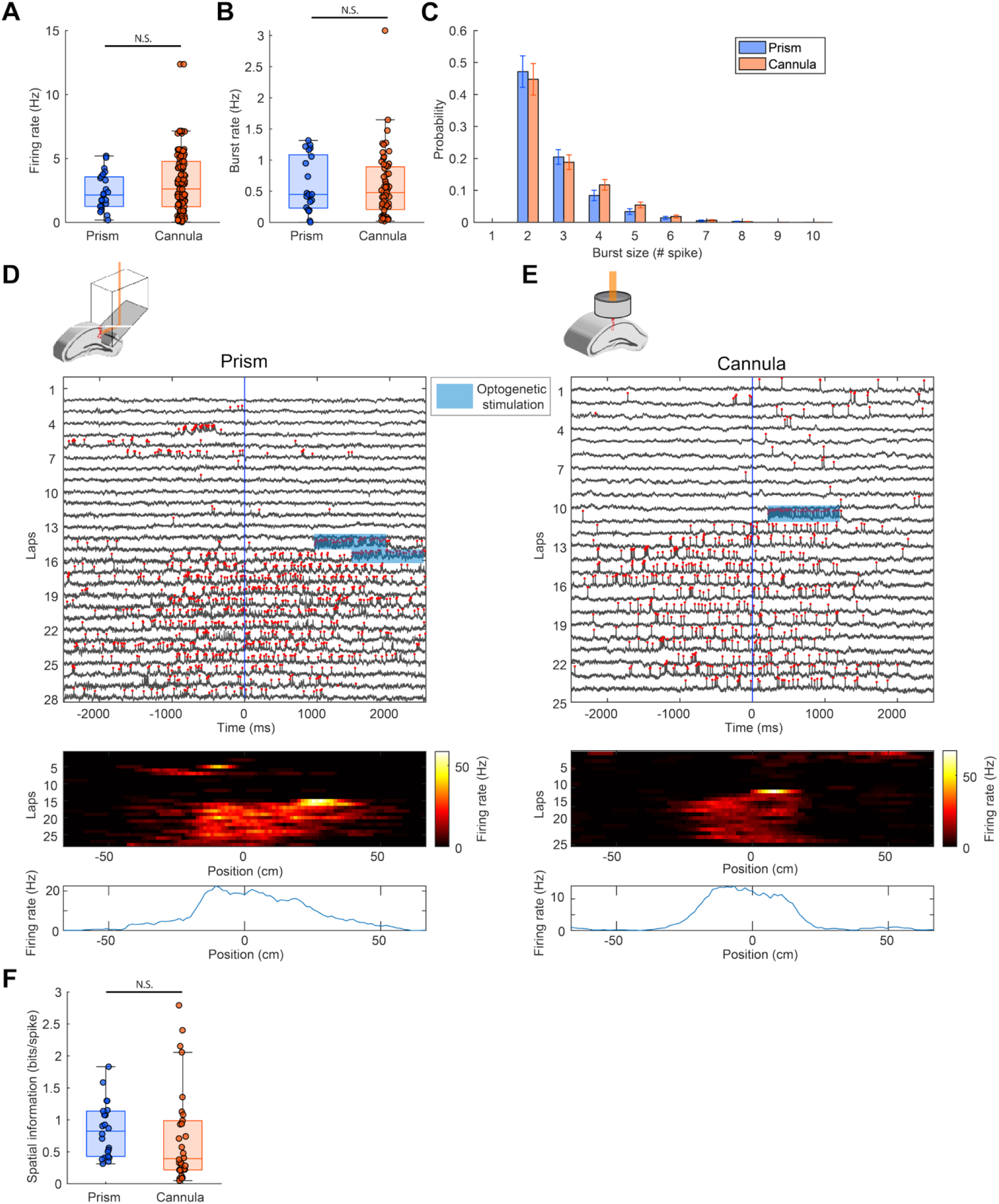
Comparison of neural firing imaged through microprism and hippocampal cannula window. Comparison of **(A)** firing rates and **(B)** burst rates of neurons from mice with microprisms (n = 22 neurons, 12 mice) and cannulas (n = 59 neurons, 3 mice). Bursts were defined as spike trains with ISI smaller than 20 ms. **(C)** Distribution of the number of spikes per burst event. **(D–E)** Examples of place field formation in neurons from mice with **(D)** microprism and **(E)** cannula. Top: voltage traces from the soma as the animal traversed the place field. Red dots indicate spikes. The blue shaded box represents optogenetic stimulation at the soma. Middle: firing maps at the place field. Following optogenetic stimulation at a specific virtual reality position, place field emerged ahead of the stimulation position. Bottom: average firing rate across laps as a function of position. **(F)** Spatial information (Methods) of neurons from mice with microprism (n = 22 neurons, 12 mice) or cannula (n = 33 neurons, 3 mice). Statistical significance was assessed with a Wilcoxon rank-sum test (N.S., P > 0.05).

**Figure S8.**
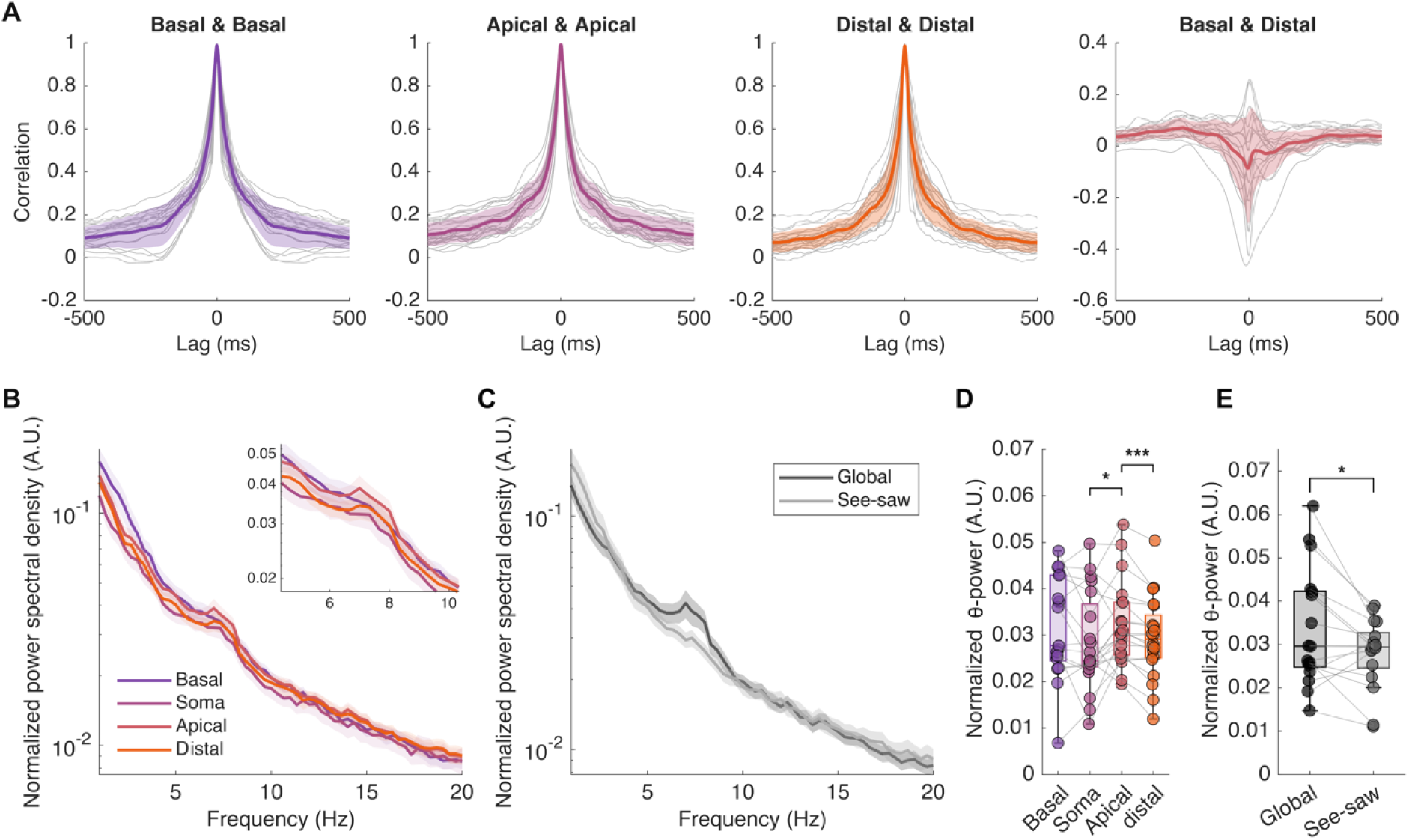
Temporal structure of dendritic subthreshold dynamics. **(A)** Auto- and cross-correlations of subthreshold voltage dynamics in different dendritic compartments. Solid line and shading indicate mean ± s.d. (*n* = 20 neurons, 12 mice). Apical distal dendrites were defined as dendritic segments branching from the main apical trunk and located ≥ 200 µm from the soma. **(B)** Power spectra of subthreshold voltages of basal dendrites, soma, apical dendrites and distal dendrites. The power spectra were normalized by the variance of the corresponding subthreshold voltage traces. Inset: zoomed view of theta-frequency power. Shading indicates s.e.m. (*n* = 20 neurons, 12 mice). **(C)** Power spectra of global and see-saw modes. **(D)** Theta-frequency (5–10 Hz) power of basal dendrites, soma, apical dendrites, and distal dendrites (*n =* 20 neurons, 12 mice; paired two-tailed t-test, \**P* < 0.05, \*\*\**P* < 0.001). **(E)** Theta-frequency power of global and see-saw modes (*n =* 20 neurons, 12 mice; paired two-tailed t-test, \**P* < 0.05).

**Figure S9.**
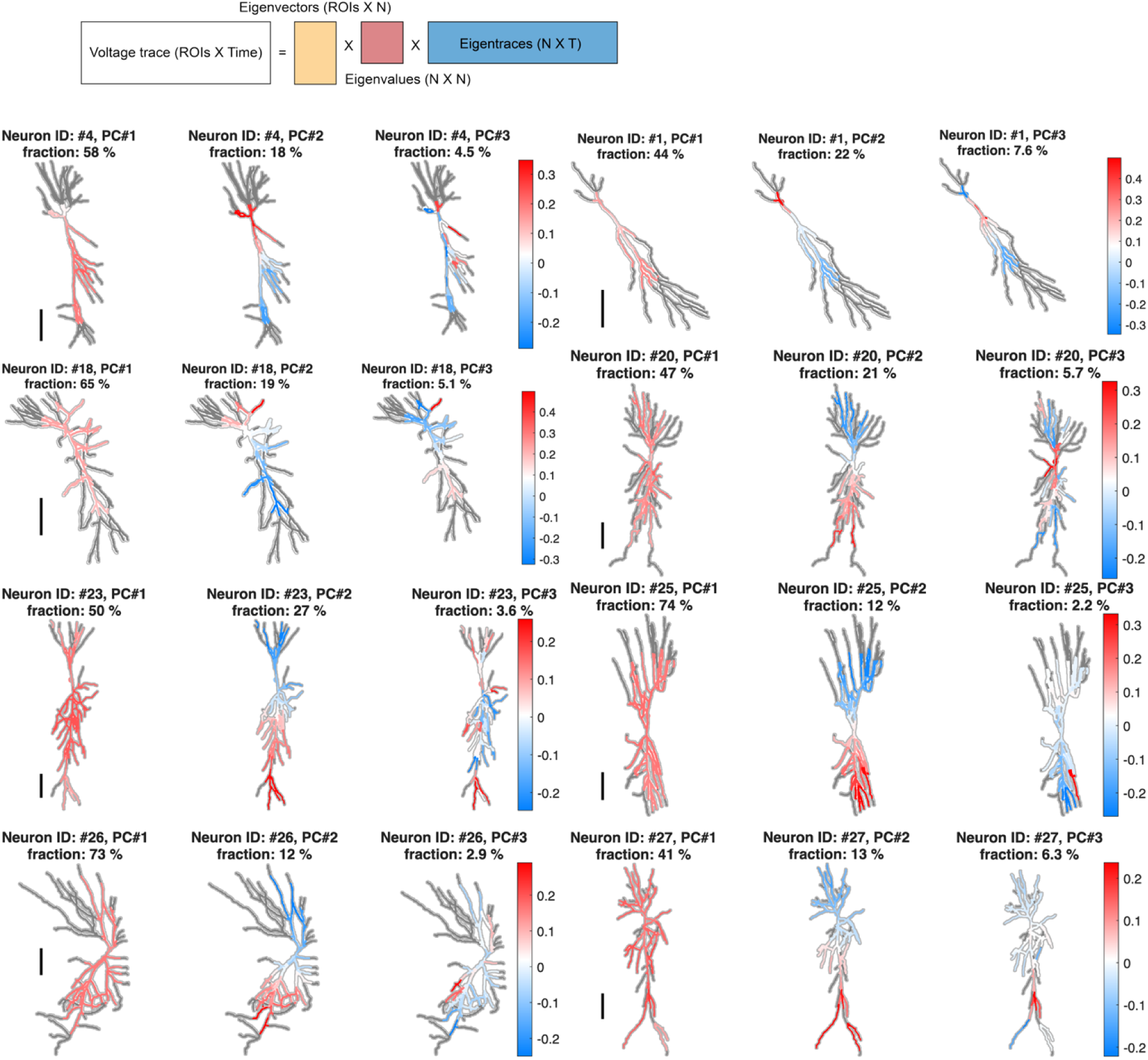
PCA of subthreshold voltage is dominated by global and see-saw modes. Eigenvectors were obtained by PCA on subthreshold voltage dynamics. The first three eigenvectors from 8 neurons are shown as examples. In all neurons, the first component was a global mode in which all dendrites fluctuated together. The second component was a see-saw mode, with basal and apical dendrites fluctuating in opposite directions. Higher-order components did not exhibit common structure across neurons and each accounted for < 10% total variance. Scale bars 100 μm.

**Figure S10.**
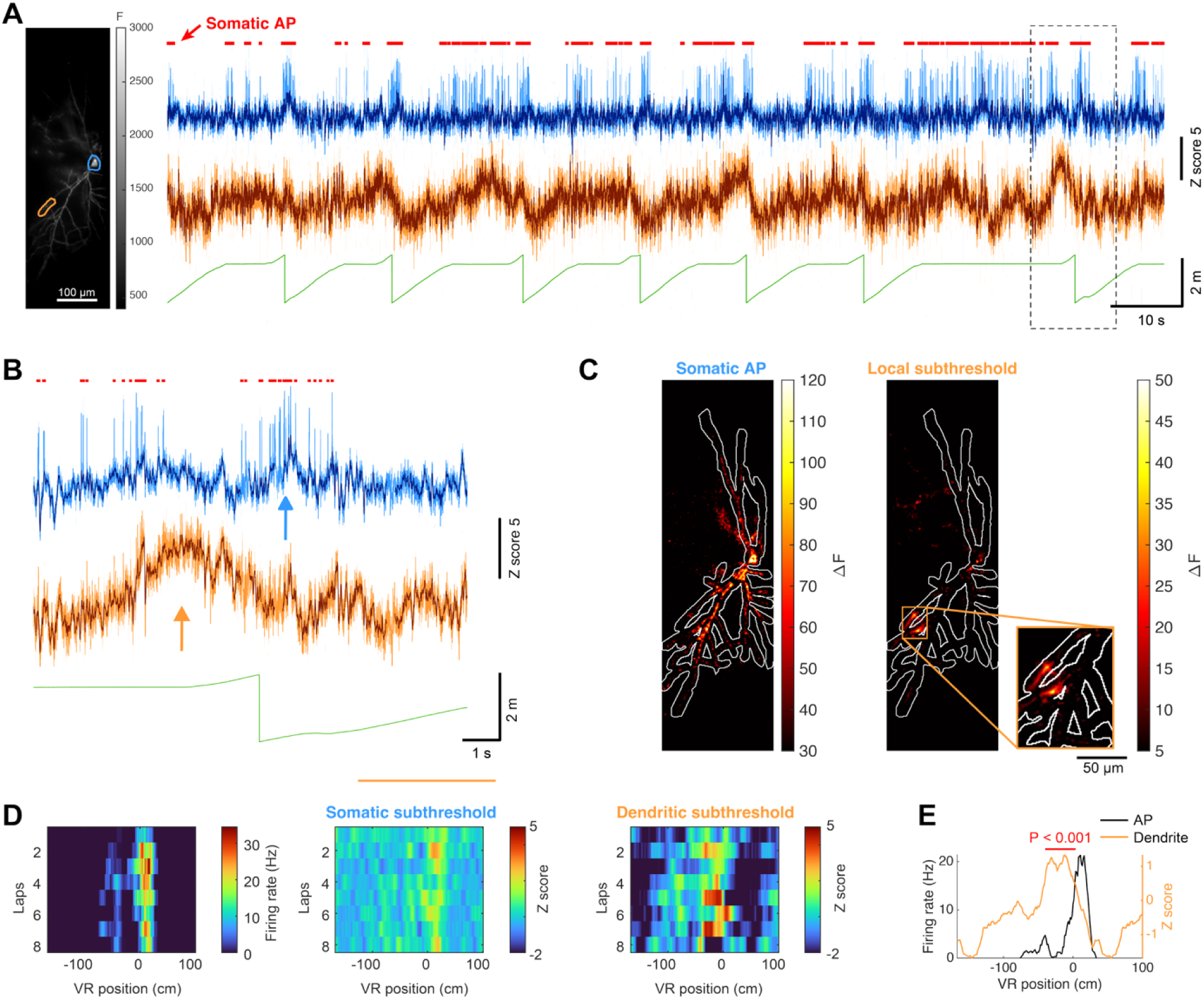
Local subthreshold events sometimes show place-field properties. **(A)** Simultaneously recorded voltage traces of soma (blue) and a dendritic segment (orange). The subthreshold voltages of soma and dendrites are shown in darker blue and orange, respectively. **(B)** Zoomed view of the dotted box in (A). The blue arrow indicates a burst of somatic action potentials, and the orange arrow marks a local subthreshold event in the dendrite. **(C)** ΔF images at the frames indicated by the blue arrow (left) and orange arrow (right) in (B). **(D)** From the left: mean firing rate, somatic subthreshold, and dendritic subthreshold as a function of VR position and lap number. **(E)** Mean firing rate (black, left axis) and dendritic subthreshold voltage (orange, right axis) as a function of VR position. The dendritic subthreshold exhibited position-tuned depolarization ahead of the place field. The red horizontal bar denotes a VR position in which the observed dendritic signal significantly exceeded the shuffled null distribution (*P* < 0.001, permutation test, 100,000 shuffles, **Methods**). Shuffling was performed by circularly shifting the voltage trace within each lap, thereby permuting position but not lap identity or temporal correlation. Events such as this were rare (only five events in two neurons, out of cumulative 10,000 s of recording from 20 neurons).

**Figure S11.**
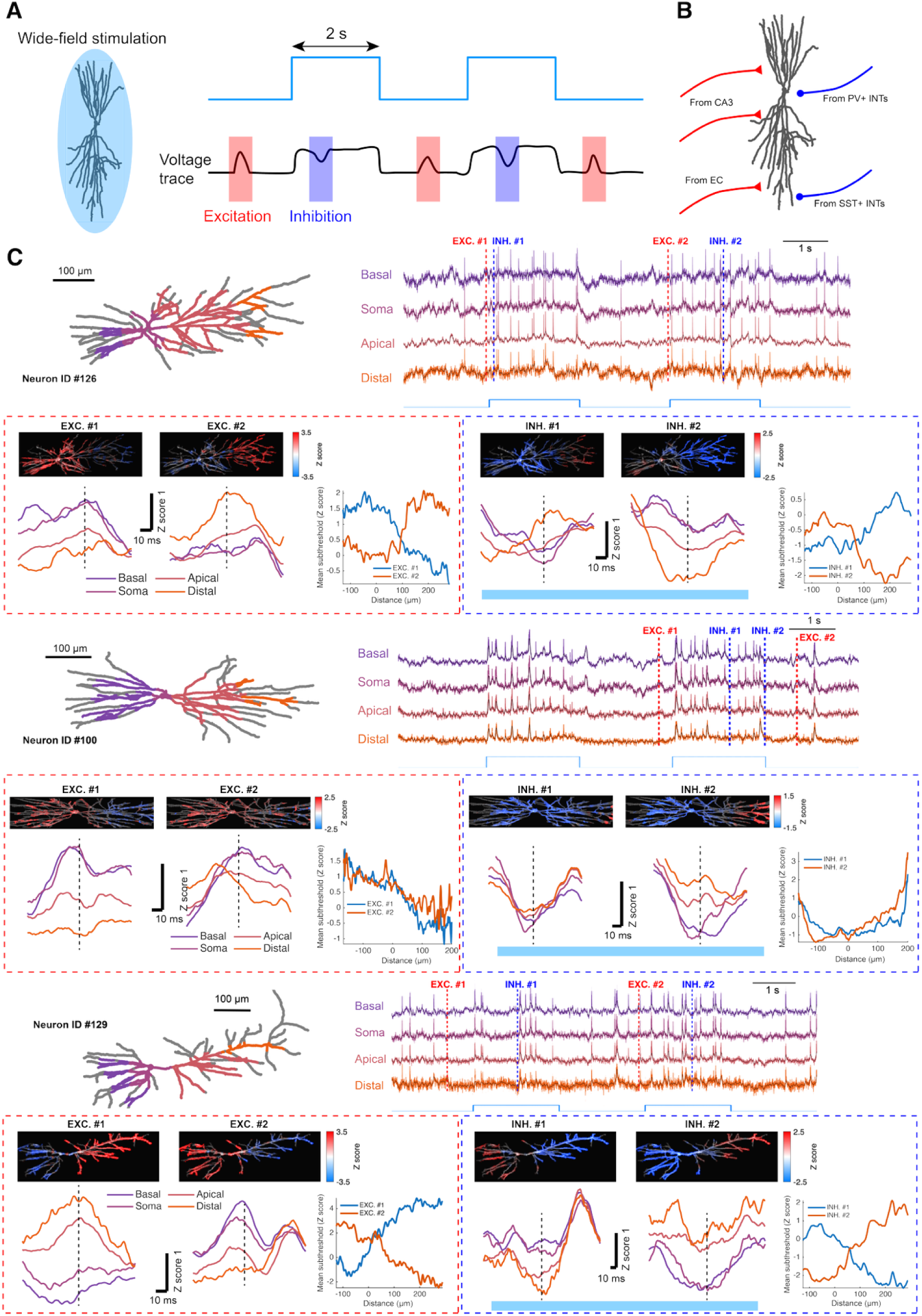
Dendritic footprints of subthreshold excitation and inhibition. **(A)** Experimental schematic. Wide-area blue light illumination was periodically applied to optically depolarize neurons, amplifying the influence of synaptic inhibition on membrane voltage. Excitatory footprints were measured during the “blue off & peak” period (red shading), and inhibitory footprints during the “blue on & dip” period (blue shading). **(B)** Major excitatory and inhibitory projections to CA1 dendrites. Excitatory inputs from CA3 and EC target proximal dendrites and distal apical dendrites, respectively. Inhibitory inputs from PV and SST interneurons project to perisomatic regions and distal apical dendrites, respectively (*35*, *38*, *46*). **(C)** Representative examples of excitatory and inhibitory footprints from three neurons. Top left: neuronal structure (grey) and voltage imaging field of view, color-coded from basal to apical dendrites. Top right: voltage traces from basal to apical dendrites. Bottom: excitatory and inhibitory footprints at the times indicated in the voltage traces. Putative excitatory footprints appeared primarily at basal and apical dendrites, whereas putative inhibitory footprints appeared in the perisomatic region and apical dendrites.

**Figure S12.**
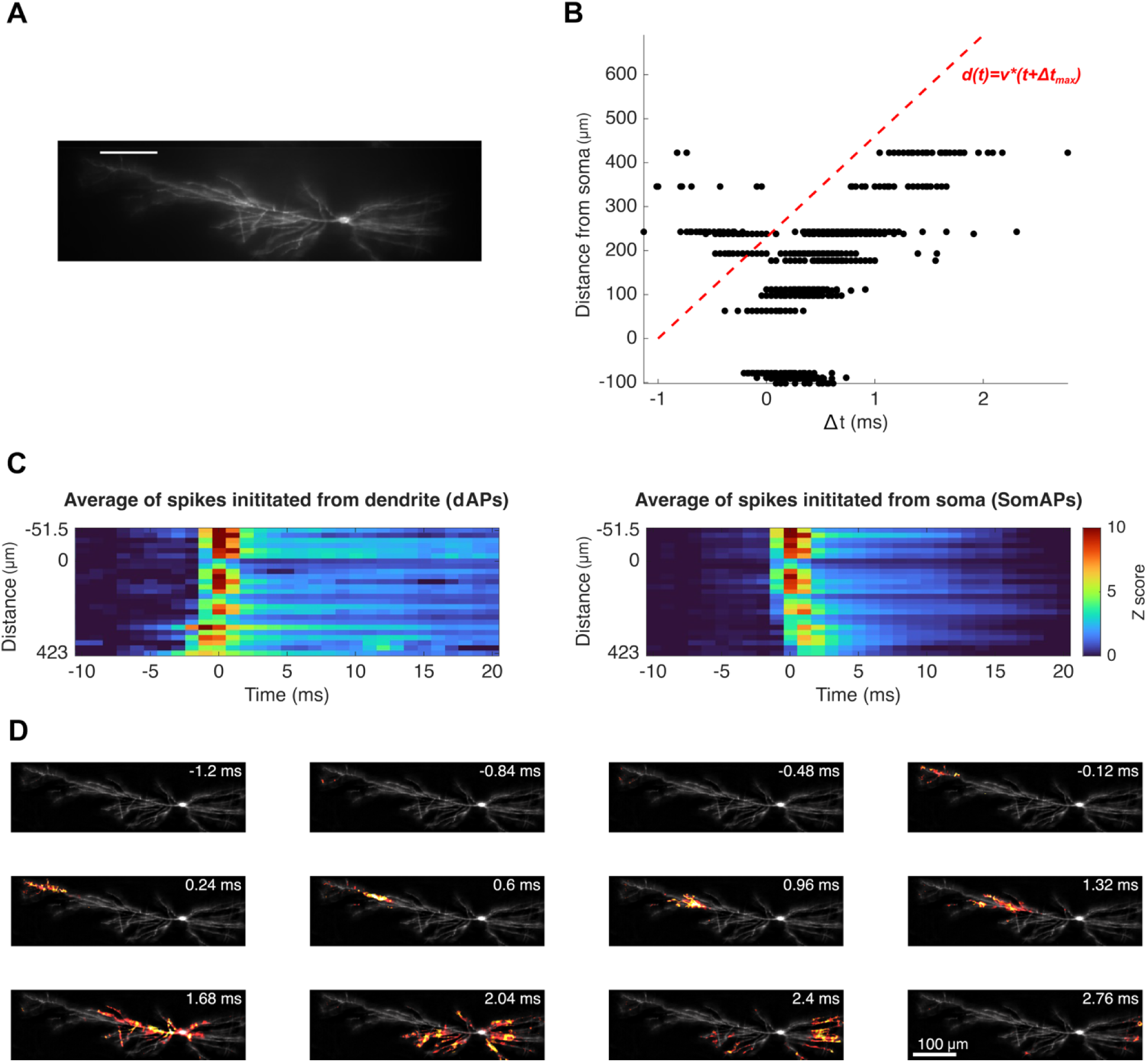
Distinguishing dAPs from bAPs. **(A)** Image of basal fluorescence (F_0_) of a neuron. Scale bar 100 μm. **(B)** Spikes detected in dendrites plotted as a function of distance from soma and the time-shift of the peak relative to the peak of the nearest somatic spike. Spline interpolation was used to estimate the spike peak timing with sub-millisecond precision. The ‘V’-shape scatter reflects a mixture of dAPs and bAPs. Red line indicates the boundary for bAPs (d(t) = v×(t+Δt_max_); v= 230 μm/ms, Δt_max =_ 1 ms (1 frame)). **(C)** Left: Average voltage kymograph triggered by dAPs (data points above red line in (B)). Right: Average voltage kymograph triggered by bAPs (data points below red line in (B)). **(D)** Snapshots from SNAPT movie showing the propagation from apical dendrite to soma of dendrite-initiated spikes. 15 out of 20 neurons exhibited dAPs at least once, and dAPs accounted for 1.0 ± 0.5% of somatic spikes (mean ± s.e.m., 101 out of 19,906 spikes, *n =* 20 neurons, 12 mice).

**Figure S13.**
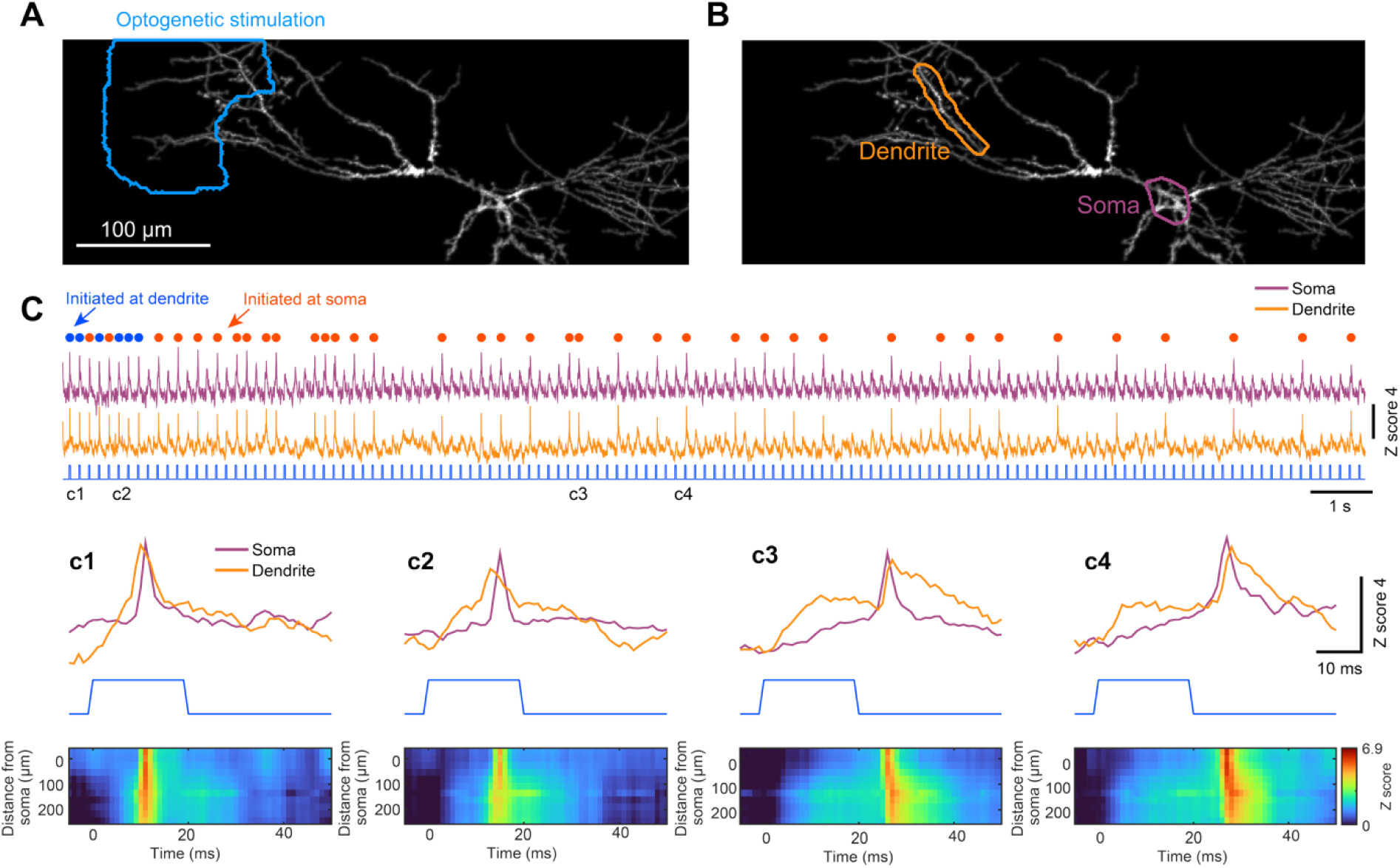
Optogenetically evoked dAPs. **(A)** Maximum z-projection of a spinning disk confocal image and blue stimulation region. **(B)** Apical (orange) and soma (purple) ROIs. **(C)** Voltage traces from the ROIs in (B) upon optogenetic pulse stimulation. Dots above the traces indicate evoked spikes (blue dots: dAPs, orange dots: soma-initiated spikes). **(D)** Four examples of optogenetically evoked spikes. c1 and c2: dAPs. c3 and c4: spikes initiated at soma leading to bAPs. Kymographs show the different directions of propagation of the dAPs vs. bAPs.

**Figure S14.**
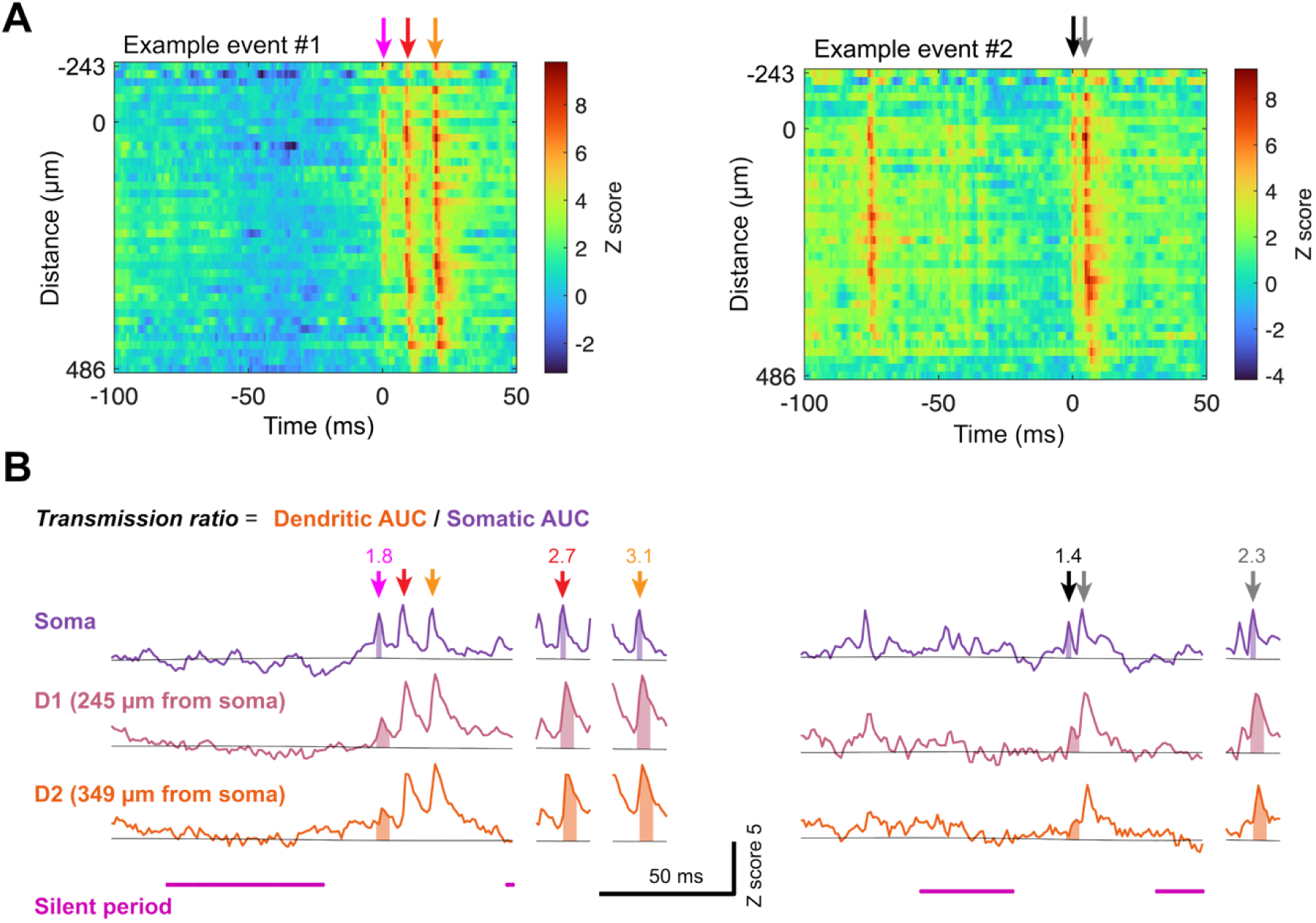
Quantifying bAP transmission. The transmission ratio was calculated as the AUC of the voltage during a bAP in apical dendrite segments > 160 μm from the soma, divided by the AUC of the spike at the soma. **(A)** Kymographs of two example events. **(B)** Voltage traces from the soma, and two dendrites for the corresponding example events in (A). Shaded regions indicate the area under the curve, and grey lines show the baseline estimated by linear interpolation between silent periods before and after the event, followed by smoothing with a 500 ms moving-average window. Numbers above the arrows indicate the transmission ratio. In both examples, the first bAP is more attenuated than the subsequent bAPs.

**Figure S15.**
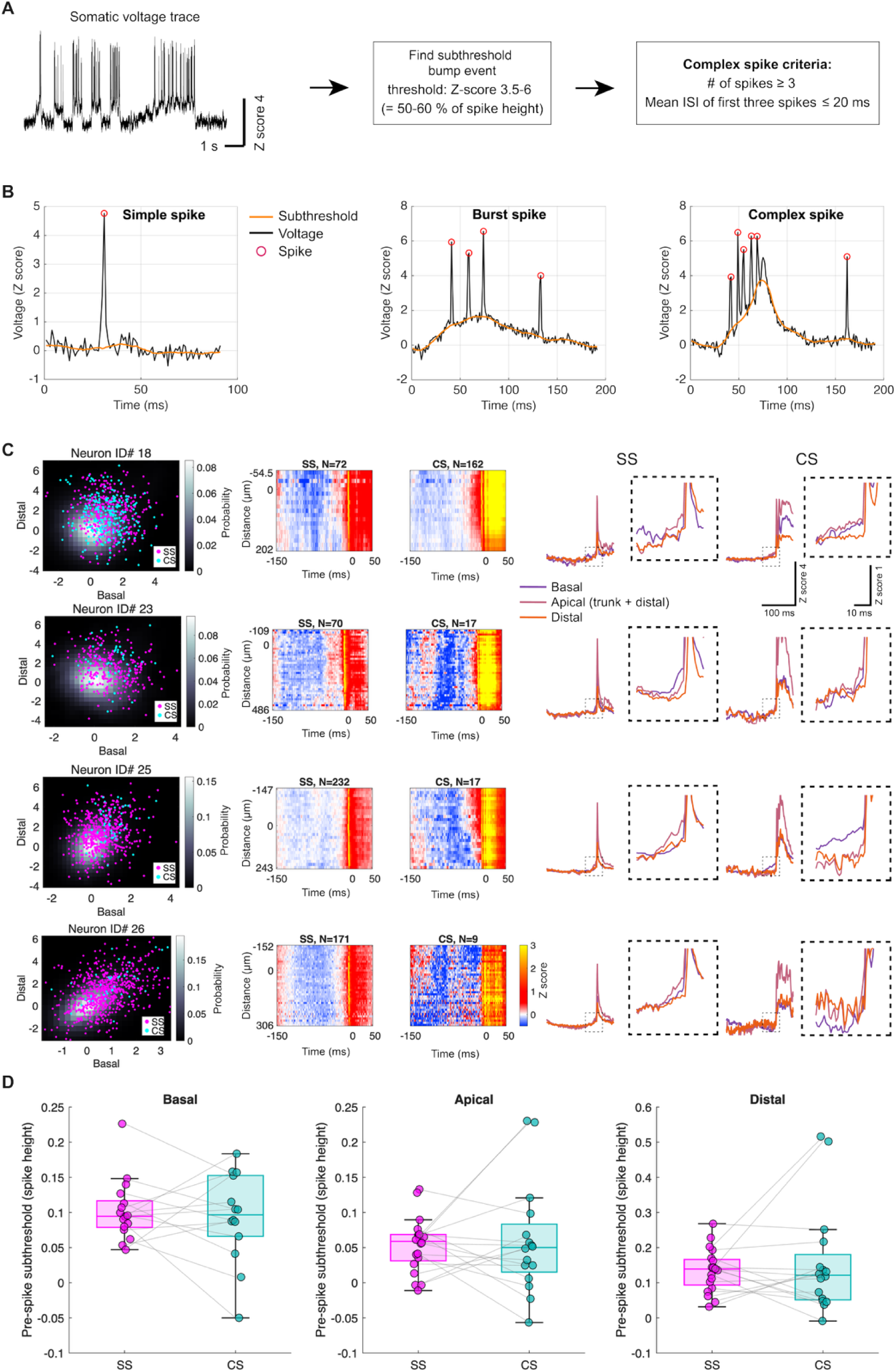
Subthreshold dynamics preceding spontaneous simple spikes vs. complex spikes. **(A)** Criteria for classifying a Complex Spike (CS). CSs contained ≥ 3 APs, with inter-spike intervals ≤ 20 ms and with a subthreshold peak amplitude exceeding a threshold. **(B)** Examples of simple, burst, and complex spike events. Bursts were defined as events with ≥ 3 APs, with inter-spike intervals ≤ 20 ms, where the subthreshold voltage did not cross the CS threshold. **(C)** Left: joint probability distributions of basal and distal subthreshold signals, across all times (grey, calculated from 1.8×10^5^–4.8×10^5^ frames), overlaid with pre-spike subthreshold of SS (magenta) and CS (cyan). Pre-spike subthreshold was defined as the mean voltage 3–10 ms before the spike. We did not observe separation of the point-clouds corresponding to SS vs. CS. Middle: Average voltage triggered by SS (left) and CS (right), shown as kymographs. Right: SS- and CS-triggered average voltage traces. **(D)** Mean pre-spike subthreshold voltage of SS and CS at basal, apical, and distal dendrites (*n =* 20 neurons, 12 mice, paired two-tailed t-test, *P* = basal: 0.63, apical: 0.63, distal: 0.44).

**Figure S16.**
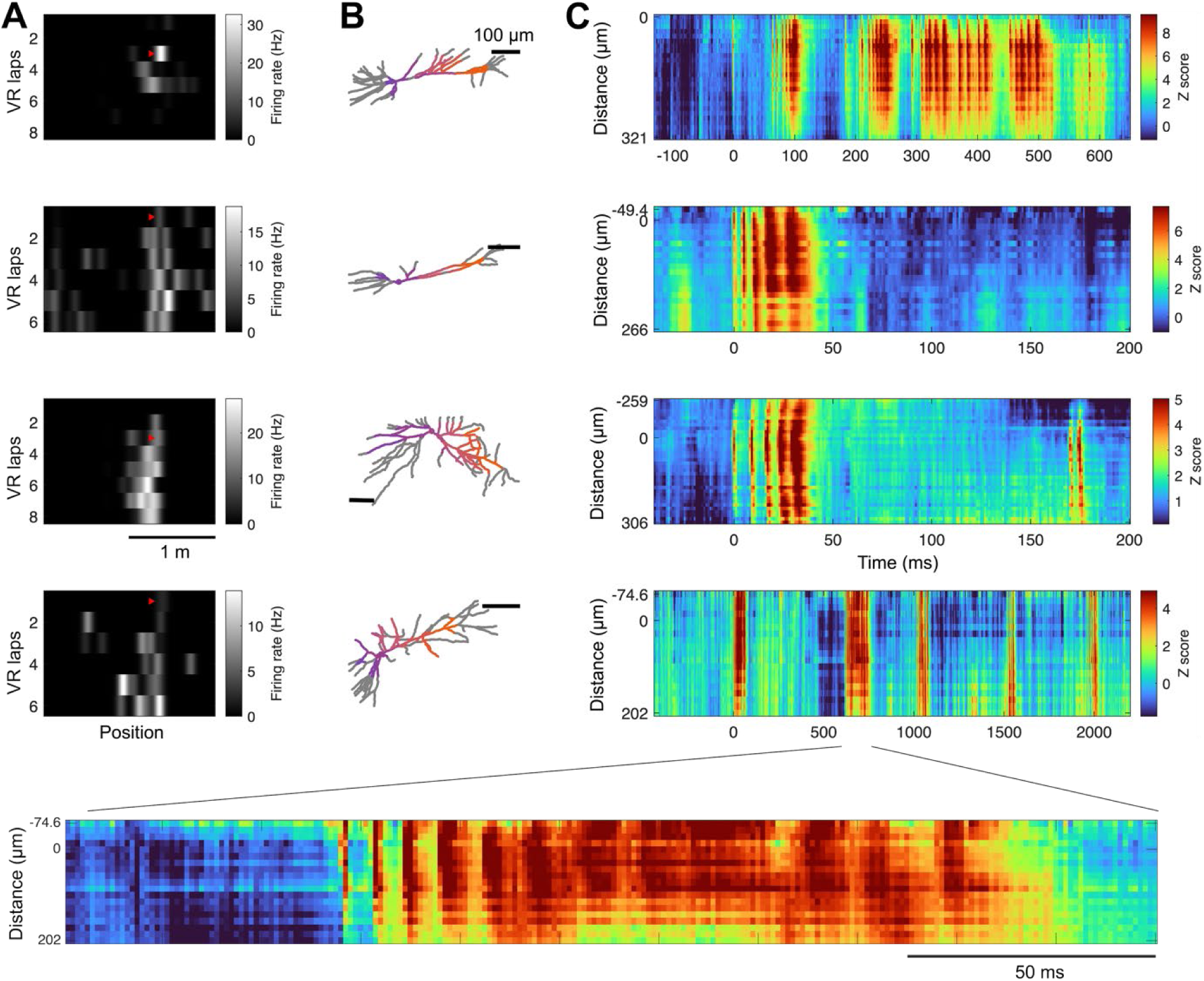
Kymographs of plateau potentials that induced place fields. Examples of place field formation and plateau potentials from four different neurons. **(A)** Mean firing rate across VR position and VR laps. The red arrows indicate the time at which the plateau potential was observed. **(B)** Neuronal morphology (grey) with the voltage-imaging field-of-view overlaid in color. Scale bars: 100 μm. **(C)** Kymographs of plateau potentials. Bottom: Magnified kymograph of the plateau potential. We did not observe distinctive subthreshold dynamics preceding BTSP-inducing plateau potentials.

**Figure S17.**
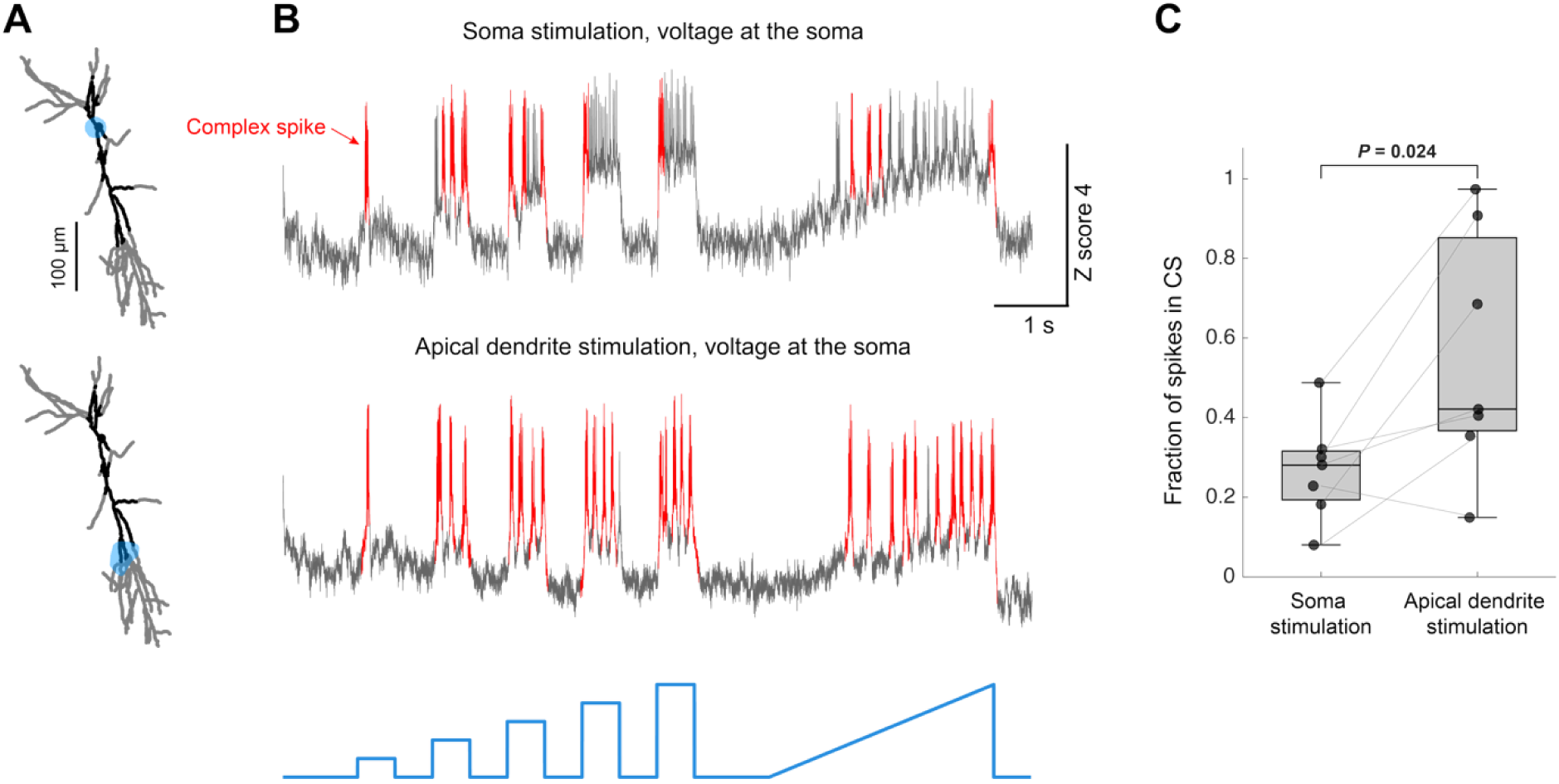
Apical stimulation favors complex spikes. **(A)** Neuronal morphology and the optogenetic stimulation region (blue shading). **(B)** Somatic voltage traces upon soma stimulation (top) and apical dendrite stimulation (bottom). Red indicates complex spikes. **(C)** Fraction of induced complex spikes during soma vs. apical dendrite stimulation (*n =* 7 neurons, 6 mice, paired two-tailed t-test, *P* = 0.024).

**Figure S18.**
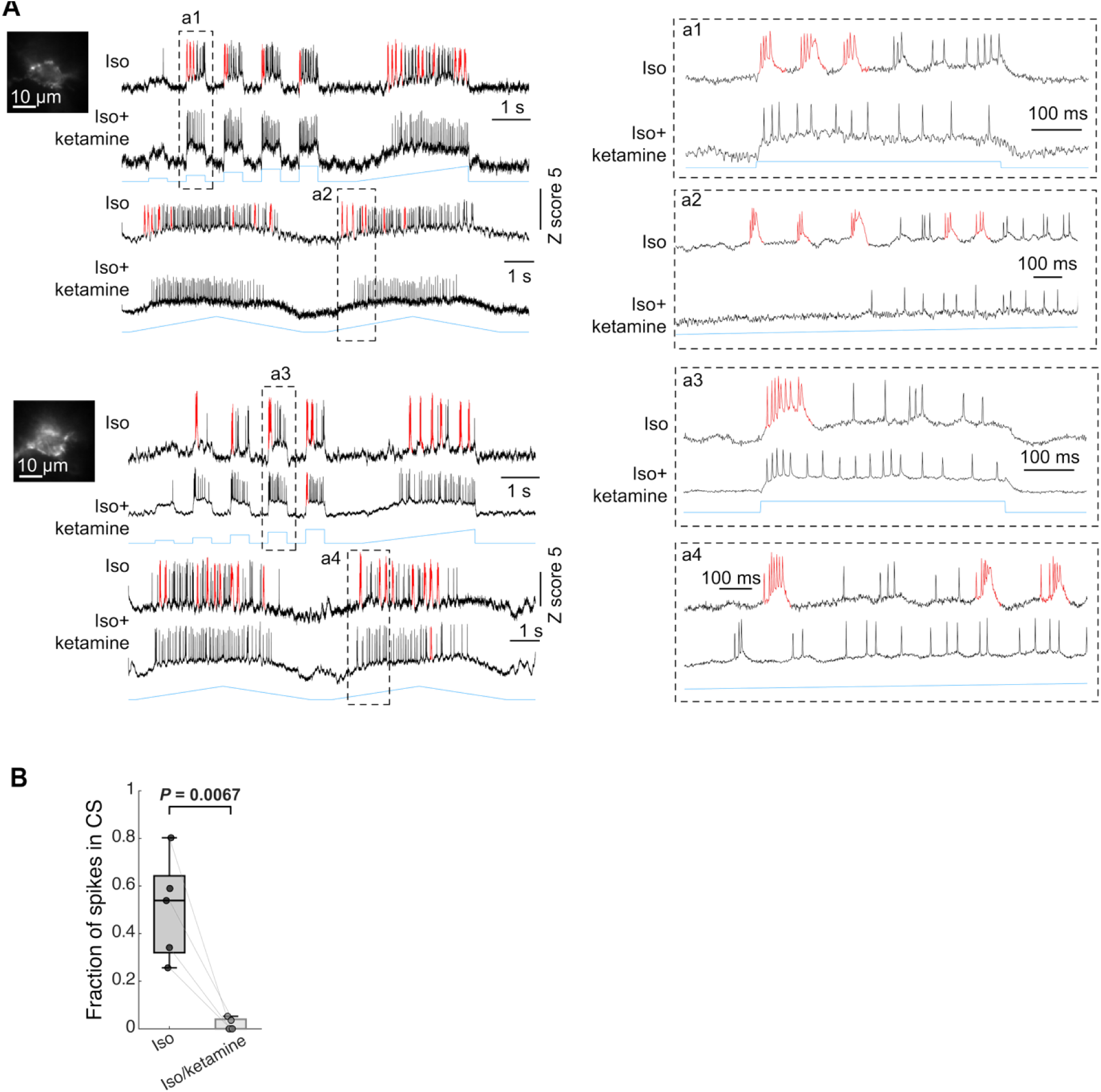
Ketamine suppresses complex spikes. **(A)** Optopatch experiment from two example neurons during isoflurane (1.5–2.5%) anesthesia and during isoflurane (1.5–2.5%) with ketamine (100 mg/kg). Ketamine was administered at least 2 hours after the isoflurane-only condition. Top: Voltage responses to step + ramp optogenetic stimulation. Bottom: Voltage responses to symmetric triangular (ramp-up and ramp-down) stimulation. Red traces denote complex spikes. **(B)** Fraction of complex spikes during isoflurane-only vs. isoflurane + ketamine (*n =* 5 neurons, 2 mice, two-tailed t-test, *P* = 0.0067).

**Figure S19.**
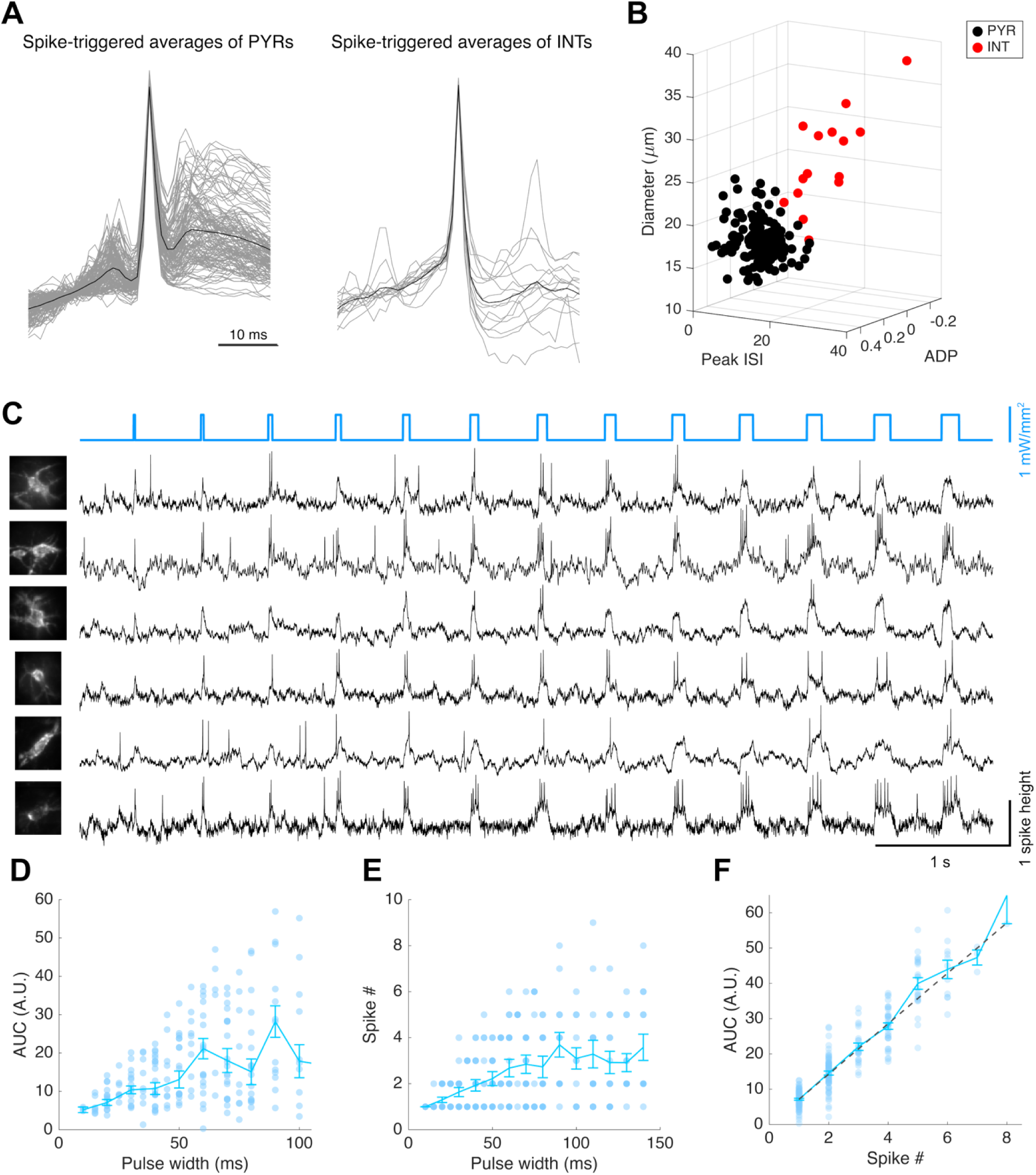
For putative interneurons, AUC scales linearly with number of spikes. **(A)** Spike-triggered average voltage waveforms of putative pyramidal neurons (PYRs, left; 28 neurons, 3 mice) and interneurons (INTs, right; 15 neurons, 2 mice). Black lines represent the population averages. **(B)** PYRs and INTs were classified based on soma diameter, peak of the distribution of inter-spike interval (ISI), and spike after-depolarization (ADP). **(C)** Same experimental protocol as in Figure 4K but applied to INTs. No CS waveforms were observed. **(D)** Cumulative AUC of evoked spikes as a function of stimulus pulse width. **(E)** Number of evoked spikes as a function of stimulus pulse width. **(F)** Relationship between cumulative event AUC and number of evoked spikes. Compared to Figure 4L (PYRs, supra-linear), INTs showed a linear increment in AUC with number of spikes. (D–F) Points show individual stimuli. Data from *n* = 1142 stimuli, 15 neurons, 2 mice.

**Figure S20.**
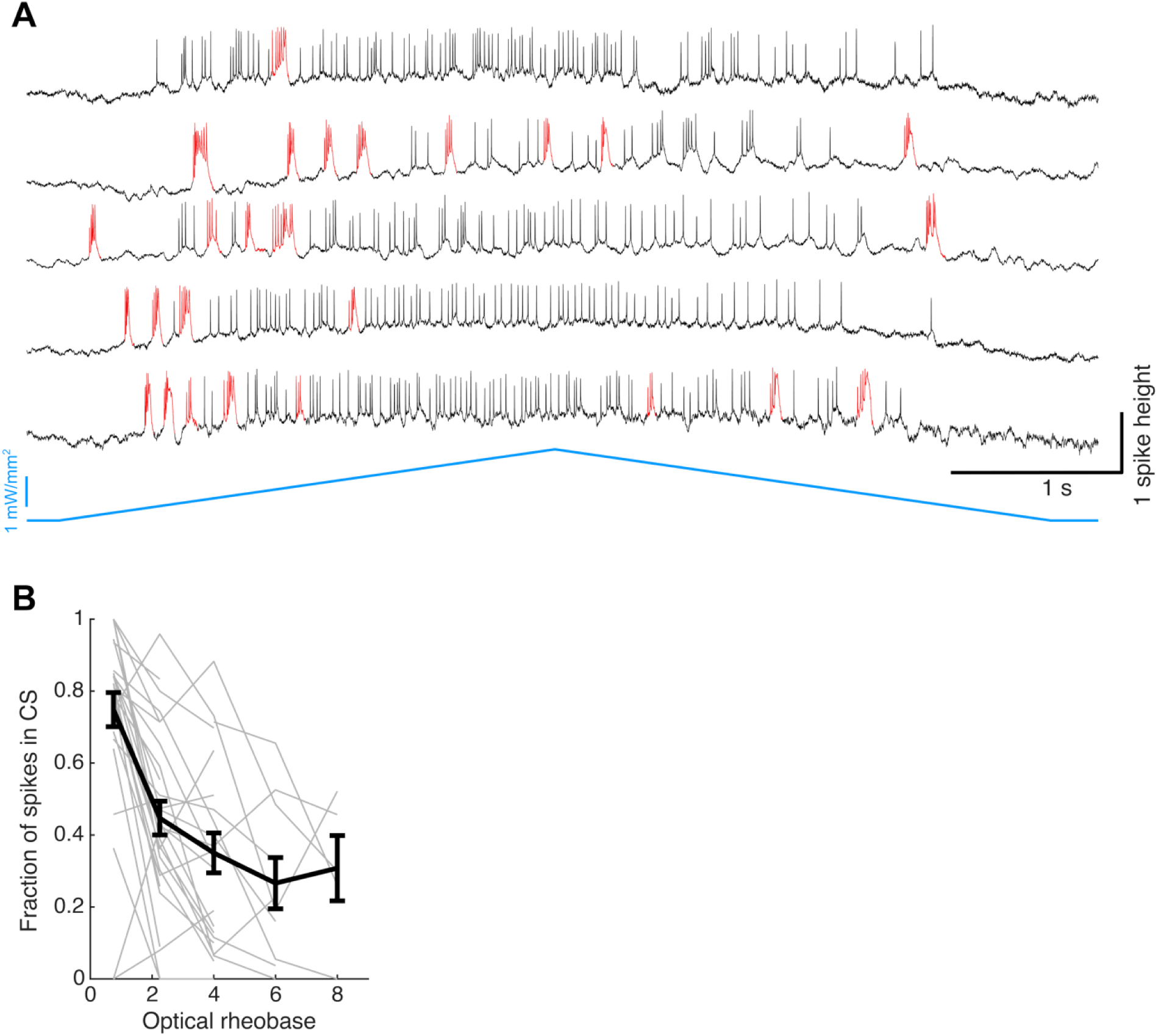
Optogenetic triangular ramp stimuli evoke CSs on rising and falling edges. **(A)** Five example somatic voltage traces from neurons in anesthetized mice. The neurons were stimulated with a triangular ramp stimulus at the soma. Red denotes CSs. **(B)** Fraction of CSs as a function of optical rheobase. Stronger blue stimulation resulted in a smaller fraction of CSs (*n* = 29 neurons, 4 mice).

